# Extracellular Matrix Physical Properties Regulate Cancer Cell Morphological Transitions in 3D Hydrogel Microtissues

**DOI:** 10.1101/2025.05.10.653220

**Authors:** Ayda Pourmostafa, Gabrielle Uskach, Mohammad Jafari, Swaprakash Yogeshwaran, Teresa L. Wood, Farid Alisafaei, Amir K. Miri

## Abstract

Cancer cells can adopt a range of morphological states linked to distinct functional behaviors during tumor progression. Some remain in a proliferative state, forming tight clusters; others detach and elongate into an invasive state; and some retain a rounded amoeboid form with minimal matrix adhesion. However, factors that determine which morphological state a cell adopts remain poorly understood. Using a combined theoretical and experimental framework, we showed that extracellular matrix (ECM) mechanics regulate cancer cell morphology in three-dimensional (3D) environments. We developed a theoretical model based on the principle of minimum energy, which predicts that a cell will adopt the morphological state—rounded, elongated, or clustered— that minimizes the total energy of the cell-ECM system. Using MDA-MB-231 breast cancer cells, we established a reliable protocol for encapsulating cells into 3D naturally-derived hydrogels with controlled stiffness and pore size. We validated the model’s predictions *in vitro* over an extended culture period. In soft ECMs, cells transitioned over time to an elongated morphology, while in stiff ECMs, cells favored clustered configurations. These transitions were governed by the physical— not chemical—properties of the hydrogel-based ECM, as confirmed by using chemically distinct yet mechanically matched composite matrices. These new insights have implications for cancer invasion modeling and potential drug screening.

## Introduction

The extracellular matrix (ECM) provides structural and mechanical cues that regulate cancer cell behavior.^1–4^ The mechanical properties of the ECM, such as stiffness and pore architecture, play crucial roles in determining how cells interact with their microenvironment.^5–9^ The mechanical properties of the native tissue can vary widely by location within the body.^10,11^ In the case of tumors, ECM remodeling leads to a wide range of mechanical properties, from soft, porous regions to dense, fibrotic areas.^12–14^ These mechanical variations influence cell adhesion, migration, and cytoskeletal organization, ultimately shaping cancer progression.^15,16^ Increased ECM stiffness, often driven by collagen deposition, is associated with tumor aggressiveness and poor patient outcomes.^17,18^ However, how cells interpret and respond to these mechanical signals in three-dimensional (3D) microenvironments remains an open question.

In traditional 2D environments, substrate stiffness is the dominant factor governing cell morphology.^19^ On a soft surface, cancer cells often retain a rounded morphology with minimal adhesion to the substrate, while on stiffer surfaces, they spread and generate higher contractile forces.^15,20^ However, in 3D ECMs, cells exhibit fundamentally different behaviors.^21–23^ Unlike in 2D, where stiffness alone dictates morphology, a synergistic combination of stiffness and pore size influences how cells respond to their surroundings in 3D ECMs.^9^ Pore size affects cell confinement, migration modes, and access to biochemical signals, making it a critical but often overlooked parameter in engineered 3D tumor models.^24–26^ This added complexity in 3D raises the fundamental question of how cells adapt their morphology when exposed to different combinations of ECM stiffness and pore size.

Understanding how cancer cells adapt their morphology within 3D microenvironments is essential for elucidating the mechanisms of tumor invasion and engineering tunable tumor models. In patient tumors, the mechanical properties of the ECM are highly heterogeneous, with regions of both soft and stiff tissue coexisting due to fibrosis, remodeling, and variations in stromal composition.^27,28^ This heterogeneity may lead cells to adopt one of three distinct morphological states: (i) a “rounded” morphology, in which the cell remains spherical and migrates through matrix pores with minimal adhesion (ameboid mode of migration); (ii) an “elongated” morphology, characterized by cell spreading and the extension of protrusions to engage with the matrix (invasive mode of migration); or (iii) a “clustered” morphology, where the cell proliferates and forms small interconnected groups.^29–32^ Each of these morphological states may have distinct implications for cell invasion and disease progression, including differences in migration potential, resistance to mechanical stress, and interaction with the tumor microenvironment.^33–35^ A quantitative framework linking ECM mechanical properties to cell morphology in 3D remains lacking in the field.

To investigate this phenomenon, we used a theoretical model based on the minimum energy principle,^36–40^ which posits that a cell will adopt the configuration that minimizes the total energy of the cell-ECM system.^41–44^ We validated these predictions experimentally by culturing individual MDA-MB-231 cells within 3D ECMs with varying mechanical properties. We used hydrogel engineering and biomaterials science to optimize the protocol for reliable 3D tumor models for up to three weeks. By integrating modeling and experimentation, our study provides new insights into the biophysical mechanisms that govern cancer cell morphology in complex microenvironments. Such insights could guide the design of new therapeutic strategies that target the physical tumor microenvironment to limit cancer invasion.

## Results

### Engineered 3D Hydrogels as a Model for Tumor ECM Mechanics

The mechanical properties of the ECM play a crucial role in shaping cell behavior in tumors, where regions of varying stiffness and pore sizes coexist due to differences in ECM density and structural remodeling. To replicate these conditions, we engineered 3D gelatin methacryloyl (GelMA) hydrogels with two distinct bulk mechanical properties: a soft ECM (~ 2 kPa) with large pores (50-100 µm) and a stiff ECM (~ 3.5 kPa) with smaller pores (10-50 µm) (**Fig. 1A–D**).^45,46^ GelMA is derived from gelatin, a denatured form of collagen, replicating native major ECM cues for tumor cells.^47^ These properties mimic the range of mechanical conditions found *in vivo*, where increased collagen concentration leads to simultaneous stiffening and pore size reduction.^17,32,48^

**Figure 1.**
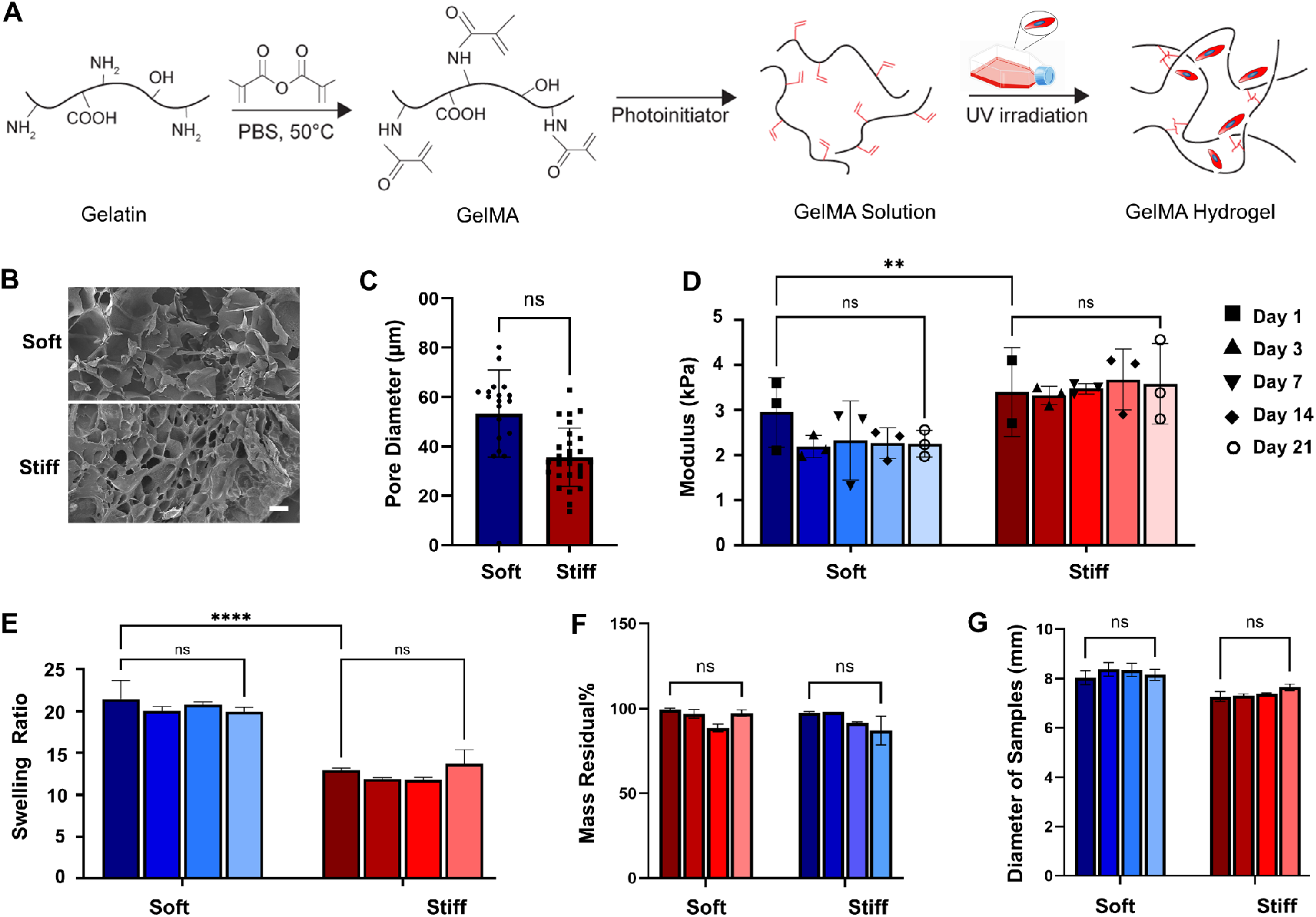
Engineering and characterization of 3D GelMA hydrogels as a stable platform for cell culture. (A) Schematic representation of hydrogel preparation and cell encapsulation. (B) Scanning electron microscopy (SEM) images of soft and stiff GelMA hydrogels (scale bar, 100 µm). (C) Quantification of pore size for soft and stiff hydrogels. (D) Stiffness measurements of acellular hydrogels over time. (E–G) Characterization of cell-laden hydrogels over 14 days, showing constant (E) swelling ratio, (F) dry mass residual percentage, and (G) hydrogel sample size, confirming system stability. ns *p* > 0.05, \**p* ≤ 0.05, \*\**p* ≤ 0.01, \*\*\**p* ≤ 0.001, \*\*\*\**p* ≤ 0.0001. Bar heights represent mean values, and error bars represent ± standard error of the mean (*n* = 4).

To assess how the mechanics of hydrogel-based ECM influence cell morphology, we encapsulated MDA-MB-231 cells within both soft and stiff ECMs (**Fig. S1-3**). In both conditions, we used a cell density of 1 × 10^6^ cells/mL, which closely mimics *in vivo* tumor settings. To ensure that the hydrogels remained structurally and chemically stable over time, we monitored swelling ratio (**Fig. 1E**), mass residual (**Fig. 1F**), and tissue diameter (**Fig. 1G**) over 14 days. All three parameters remained constant, indicating that the hydrogel structure was preserved throughout the experiment. In particular, the mass residual remained unchanged, suggesting minimal degradation and confirming that matrix metalloproteinase (MMP) activity did not significantly alter the ECM. These results demonstrate that our hydrogel system provides a stable and physiologically relevant platform to study how ECM mechanics regulate cell morphology, as suggested by our previous study.^26^

### Energy-Based Model Predicts Elongated Morphology in Soft ECM and Clustered Morphology in Stiff ECM

To predict which morphological state (rounded, elongated, or clustered) a cell is most likely to adopt in the ECM, we used a theoretical model based on the minimum energy principle. Based on this principle, we hypothesized that a cell will take the shape that requires the least total energy. We implemented the model within a three-dimensional finite element framework to simulate cell spreading and contraction across three morphological states: rounded, elongated, and clustered (see **Materials and Methods** and **SI 1-3**). For each morphological state, we calculated the key energy components contributing to the total cellular energy in both soft and stiff extracellular matrices (**Fig. 2A, S4**). In this framework, a positive energy component indicates resistance to a particular morphology, with higher values representing greater opposition to that state. Conversely, a negative energy component favors a given morphology, with more negative values indicating a stronger preference for that state.

**Figure 2.**
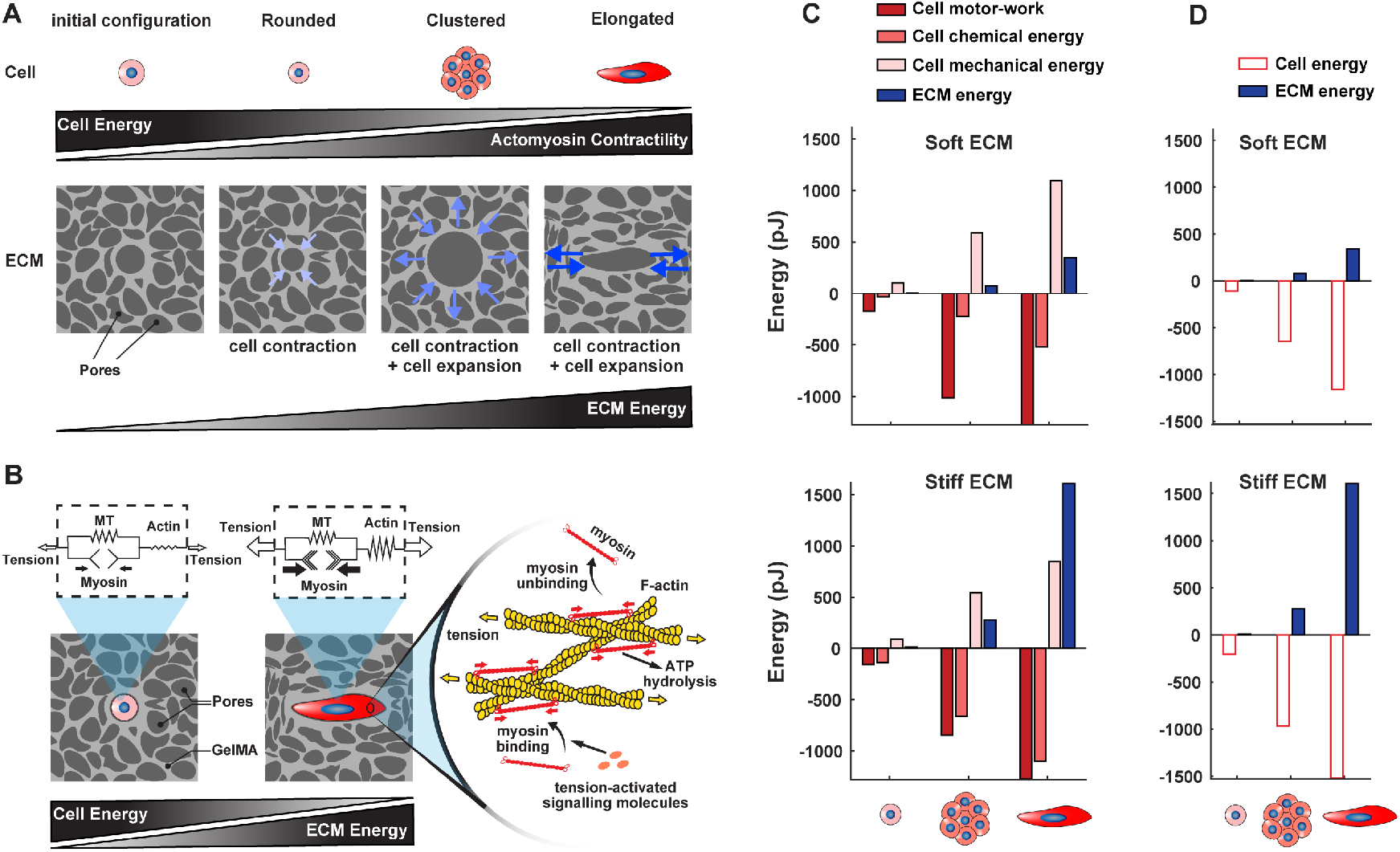
Energy-based model predicts distinct energy profiles for rounded, elongated, and clustered morphologies. (A–B) Cells initially exhibit a rounded morphology within the extracellular matrix (ECM). Over time, they may either maintain this rounded shape or transition into elongated or clustered morphologies. The model predicts that the elongated morphology has the highest ECM mechanical energy because elongation requires substantial deformation of the surrounding ECM. However, elongated cells have the lowest cell energy due to elevated actomyosin contractility, which results from ATP hydrolysis and the associated release of chemical energy. This chemical energy release is the primary factor lowering the total cell energy in elongated cells. (C) Detailed breakdown of energy components: Cell energy was estimated as the sum of three key contributions — (i) chemical energy released during ATP hydrolysis as myosin motors bind to actin filaments, (ii) motor-work energy, representing the mechanical work generated by actomyosin contractility, and (iii) cell mechanical energy, which includes the passive strain energy stored in cytoskeletal structures and the external work performed by them. Each of these three energy components and the ECM mechanical energy was estimated and plotted. Positive energy values indicate resistance to adopting a given morphology, while negative values favor it. (D) The three cell energy components were summed to obtain the total cell energy, which was plotted alongside ECM mechanical energy. The results show that the elongated morphology has the lowest total cell energy but the highest ECM mechanical energy, while the rounded morphology has the lowest ECM mechanical energy but the highest cell energy. The clustered morphology lies between these two extremes.

One major component of the total energy is ECM mechanical energy, which represents the energy a cell must expend to deform the ECM in order to take a particular morphology. Initially, cells are small and round, measuring approximately ~ 20 µm in diameter, as shown in our experiments in **Fig. S5**. For a cell to transition into a mesenchymal-like morphology, it must spread and push against the surrounding ECM, elongating its long axis to roughly 100 µm, as observed experimentally in **Fig. S5**. This process requires mechanical work, making ECM mechanical energy an unfavorable factor for elongated morphology. However, for a rounded morphology, ECM mechanical energy remains negligible because the initial cell size (~ 20 µm) is equal to or smaller than the ECM pore size (measured in **Fig. 1C**). This trend is schematically illustrated in **Fig. 2A**, where the rounded state has the lowest ECM mechanical energy, while the elongated state has the highest. Therefore, based solely on ECM mechanical energy, the rounded morphology should be the most favorable.

However, total energy also includes cell energy, which we estimate as the sum of three key components (**SI 2.1**): (i) chemical energy, which is released when myosin motors bind to actin filaments, hydrolyzing ATP; (ii) motor-work energy, which represents the mechanical work performed by actomyosin contractility; and (iii) cell mechanical energy, which includes both the passive strain energy stored in cytoskeletal components (such as microtubules and actin filaments) and the external work performed by these structures. We estimated each of these three components separately (**Fig. 2C**), and then summed them to obtain the cell energy. This cell energy was plotted alongside the ECM energy in **Fig. 2D**.

These energy components vary depending on cell morphology, since different shapes require different levels of contractile machinery activation and cytoskeletal organization. We recently demonstrated that cell elongation significantly increases actomyosin-based contractility, resulting in enhanced phosphorylation of myosin and greater actin polymerization.^39^ This, in turn, increases ATP hydrolysis and chemical energy release. As a result, our model predicts that the elongated morphology exhibits the highest actomyosin contractility (**Fig. 2A,B**), yielding the most negative cell energy values (**Fig. 2D**). Conversely, the rounded morphology is associated with the lowest contractility and, therefore, the least negative cell energy. In summary, the elongated state has the lowest cell energy but the highest ECM mechanical energy, while the rounded state exhibits the opposite trend. The clustered state falls between these two extremes (**Fig. 2A**).

By summing ECM mechanical energy and cell energy, we calculated the total energy for each morphological state in both soft and stiff ECMs (**Fig. 3A**). The model predicted that in soft ECM, the elongated morphology has the lowest total energy, making it the most favorable state, followed by the clustered and rounded morphologies. In contrast, in stiff ECM, the clustered morphology has the lowest total energy, followed by the rounded morphology, while the elongated morphology is the least favorable. This prediction arises because, in soft ECM, the energy cost of spreading and elongating is relatively low due to the ECM’s low stiffness and large pore size (**Fig. 2D**). Subsequently, the cell can afford this mechanical cost in exchange for achieving high actomyosin contractility, which is energetically beneficial by releasing chemical energy. However, in stiff ECM, the energy required to deform the ECM is significantly higher, making elongation energetically prohibitive. As a result, clustered and rounded morphologies become more favorable in stiff ECM conditions (**Fig. 3A**).

**Figure 3.**
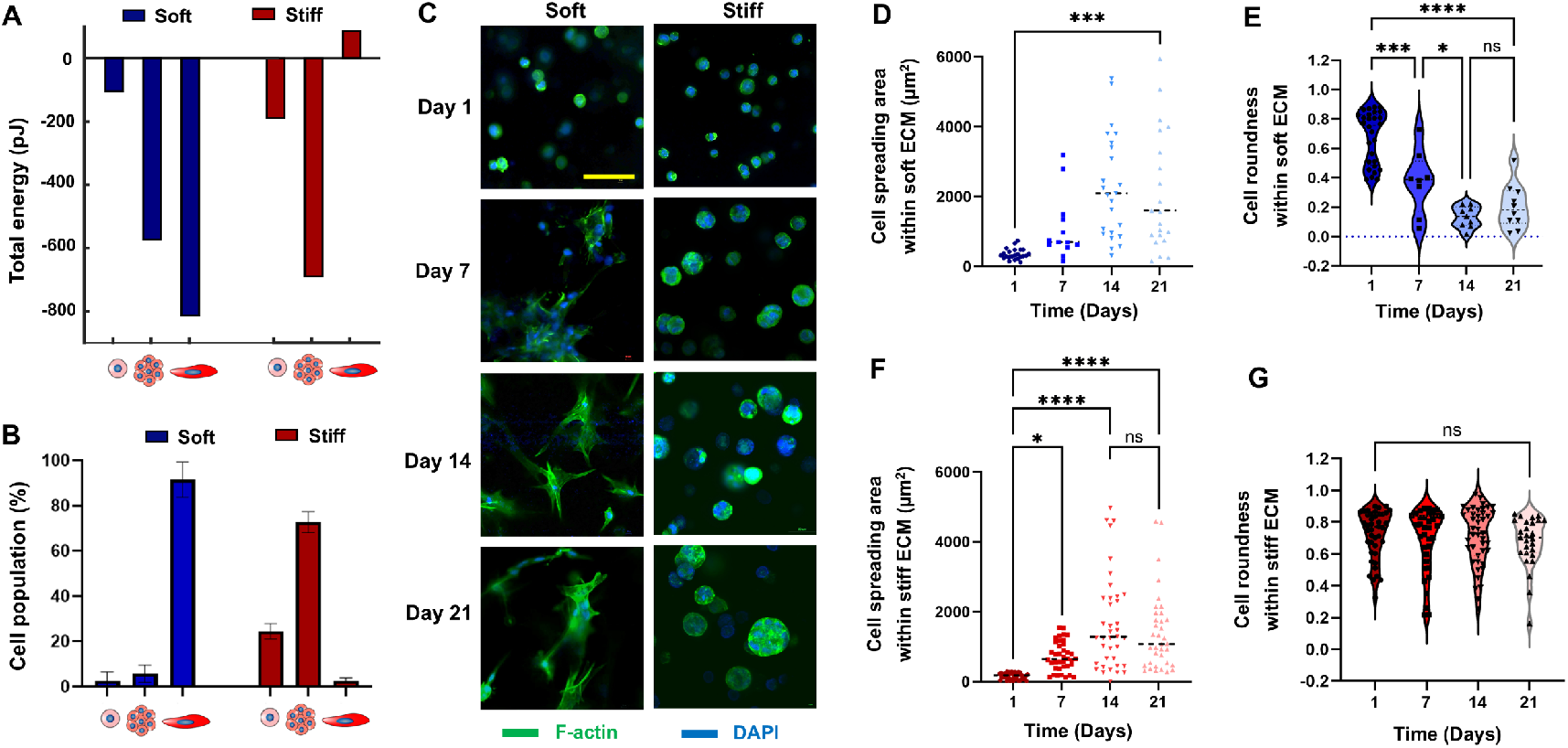
Experimental validation of model predictions: ECM stiffness dictates distinct morphological transitions. (A) Total energy for each morphological state was estimated by summing ECM mechanical energy and cell energy in both soft and stiff ECMs. The model predicts that in soft ECM, the elongated morphology has the lowest total energy, making it the most favorable state, followed by clustered and rounded morphologies. In stiff ECM, the clustered morphology has the lowest total energy, followed by the rounded morphology, while the elongated morphology is the least favorable. (B) Experimental validation of model predictions by quantifying the percentage of cells adopting different morphological states (rounded, clustered, elongated) in soft and stiff ECMs. (C) Representative images illustrating morphological changes of cells within soft and stiff ECMs over a 21-day period, visualized by immunofluorescence staining of F-actin (green, phalloidin) and nuclei (blue, DAPI). (scale bar, 50 µm). Images are from three independent biological replicates, each containing three hydrogel technical replicates and triplicate fields of view per sample. (D–G) Quantification of cell area and shape evolution in soft and stiff ECMs over 21 days. At day 1, cells exhibited similar rounded morphologies with low spreading areas in both ECM types. Over time, cells in soft ECM adopted an elongated morphology, as indicated by a gradual reduction in roundness, whereas cells in stiff ECM proliferated and formed clusters while maintaining a rounded shape. ns *p* > 0.05, \**p* ≤ 0.05, \*\**p* ≤ 0.01, \*\*\**p* ≤ 0.001, \*\*\*\**p* ≤ 0.0001. Bar heights represent mean values and error bars represent ± standard error of the mean (*n* = 4).

### Validation of Model Predictions: ECM Mechanical Properties Dictate Distinct Cell Morphological Transitions

To test the model predictions, we cultured MDA-MB-231 cells within both soft and stiff ECMs and monitored their morphological transitions over 21 days. Initially, at day 1, cells in both soft and stiff ECMs predominantly exhibited a rounded morphology (**Fig. 3C**). However, over time, distinct transitions emerged depending on ECM stiffness. In soft ECM, cells gradually transitioned from individual rounded morphologies to an elongated state, whereas in stiff ECM, they transitioned from a rounded morphology to a clustered state.

We quantified these morphological changes in **Fig. 3B**, which presents the percentage of cells in each state (rounded, elongated, and clustered) across different ECM conditions. Consistent with model predictions (**Fig. 3A**), elongated cells became the dominant population in soft ECM, while clustered cells were the most prevalent in stiff ECM.

Further quantitative analysis is shown in **Fig. 3D-G**, where we tracked the evolution of cell shape over time. At day 1, cells in both ECM conditions exhibited similar round morphologies with low spreading areas (**Fig. 3D,F**). However, in soft ECM, cells progressively adopted an elongated morphology over 21 days, as evidenced by a gradual reduction in roundness (**Fig. 3E**). In contrast, in stiff ECM, cells proliferated, forming clusters while maintaining their spherical shape, reflected by their consistent roundness over time (**Fig. 3G**). Our experimental results validate the model predictions, demonstrating that ECM stiffness governs distinct morphological transitions: elongation in soft ECM and clustering in stiff ECM.

### ECM Mechanical Properties, Not Chemical Composition, Govern Cell Morphological Transitions

To determine whether the observed cell morphological transitions were dictated by ECM mechanical properties rather than chemical composition, we prepared two new ECMs composed of a hybrid hyaluronic acid methacrylate (HAMA)-GelMA hydrogel, in contrast to the GelMA-only ECMs used previously. While these composite ECMs differed chemically from GelMA ECMs, our measurements confirmed that they maintained similar mechanical properties. Specifically, similar to the GelMA ECMs in **Fig. 1**, the composite ECMs exhibited distinct mechanical properties: a soft ECM (~ 0.5 kPa) with large pores and a stiff ECM (~ 3.5 kPa) with smaller pores (**Fig. 4A-C, S6**). Additionally, swelling ratio (**Fig. 4D**) and mass residual (**Fig. 4E**) measurements for the composite ECMs closely matched those of the GelMA ECMs (**Fig. 1E-F**). Notably, the mass residual remained unchanged, indicating minimal ECM degradation and confirming that cell produced-MMP did not significantly alter the ECM.

**Figure 4.**
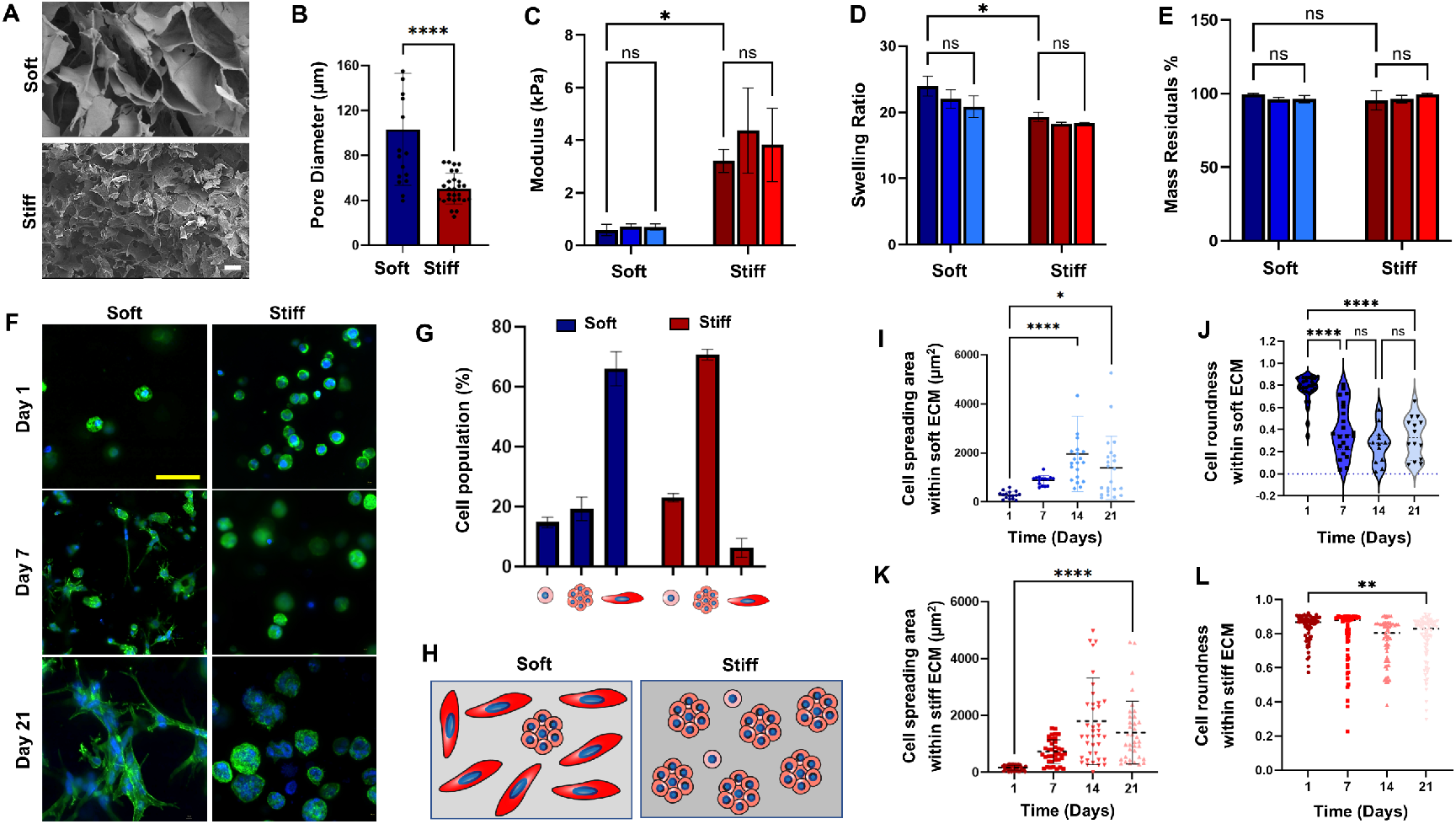
Composite ECMs replicate mechanical regulation of cell morphology independent of chemical composition. (A) SEM images of soft and stiff composite HAMA-GelMA hydrogels (scale bar, 100 µm). (B) Quantification of pore size in soft and stiff composite hydrogels. (C–E) Characterization of mechanical and structural stability of composite hydrogels over time, including (C) compressive modulus, (D) swelling ratio, and (E) mass residual of cell-laden hydrogels measured at days 1, 7, and 21. (F) Representative immunofluorescence images of F-actin (green, phalloidin) and nuclei (blue, DAPI) showing cell morphology in composite hydrogels (scale bar, 50 µm). Images are from three independent biological replicates, each containing three hydrogel technical replicates and triplicate fields of view per sample. (G) Quantification of the percentage of cells adopting different morphological states (rounded, clustered, elongated) in soft and stiff composite hydrogels. (H) Schematic illustration of cell morphology within soft and stiff composite hydrogels. (I–L) Quantification of cell area and shape evolution in soft and stiff composite hydrogels over 21 days. ns *p* > 0.05, \**p* ≤ 0.05, \*\**p* ≤ 0.01, \*\*\**p* ≤ 0.001, \*\*\*\**p* ≤ 0.0001. Bar heights represent mean values, and error bars represent ± standard error of the mean (*n* = 4).

After verifying that the composite ECMs replicated the physical properties of the GelMA ECMs, we examined whether differences in chemical composition influenced cell morphology. If ECM chemistry played a significant role, we would expect differences in cell morphological transitions between composite and GelMA ECMs. However, our results showed that cell behavior remained consistent across both ECM types. Similar to the GelMA ECMs, at day 1, cells in both soft and stiff composite ECMs exhibited a rounded morphology (**Fig. 4F**). Over time, distinct transitions emerged depending on ECM stiffness, mirroring our previous results in **Fig. 3**. In the soft composite ECM, cells gradually transitioned from individual rounded morphologies to an elongated state, whereas in the stiff composite ECM, they shifted toward a clustered state (**Fig. 4F**).

Quantification of these morphological changes further confirmed the consistency of results between ECM types. As observed in GelMA ECMs, elongated cells became the dominant population in soft composite ECMs, while clustered cells were most prevalent in stiff composite ECMs (**Fig. 4G,H**). Tracking the evolution of cell shape over time showed that, at day 1, cells in both composite ECMs displayed rounded morphologies with low spreading areas (**Fig. 4I,K**). In soft composite ECMs, cells progressively adopted an elongated morphology over 21 days, as reflected by a gradual reduction in roundness (**Fig. 4J**). In contrast, cells in stiff composite ECMs proliferated, forming clusters while maintaining their spherical shape, as indicated by their consistent roundness over time (**Fig. 4L**). Overall, these findings reinforce our results from GelMA ECMs and validate our model predictions, demonstrating that ECM physical properties—rather than chemical composition—govern cell morphological transitions.

## Discussion and Conclusions

In this study, we combined theoretical modeling and experimental validation to uncover how the physical properties of the ECM guide cancer cell morphology in three-dimensional environments. By balancing the mechanical energy cost of ECM deformation against the energetic gains from actomyosin contractility, our model predicted that cells would adopt the morphological state that minimizes total energy. These predictions were confirmed experimentally, demonstrating that ECM physical properties dictate whether individual cancer cells become rounded, elongated, or clustered within 3D matrices.

A key insight emerging from our work is that different morphological states likely represent distinct functional programs with important implications for tumor invasion. Cells that adopt an elongated morphology exhibit high levels of actomyosin contractility, a hallmark of migratory and invasive behavior. In contrast, cells that form clusters prioritize proliferation and cell-cell adhesion, promoting tumor expansion. Thus, ECM mechanics may not only determine cell shape but also bias the balance between two fundamental cancer strategies: local tumor growth versus systemic dissemination through invasion.

This duality reveals an important principle: softer and more porous ECM environments may prime cancer cells toward an invasive, migratory phenotype, whereas stiffer and denser environments may favor clustering and tumor mass expansion. Given that patient tumors are mechanically heterogeneous, with regions of variable stiffness and porosity, this mechanical heterogeneity could locally program cancer cells to either grow in place or escape and invade surrounding tissues.

Together, our theoretical and experimental findings offer a new biophysical perspective on how cancer cells adapt to heterogeneous extracellular matrix environments. Future work may extend these principles to other cell types and investigate therapeutic approaches to manipulating the mechanical landscape of tumors to design, control, and monitor cancer cell behavior.

## Materials and Methods

### Cell Preparation

A human breast cancer cell line (MDA-MB-231) was purchased from ATCC and was cultured in Dulbecco’s Modified Eagle Medium mixed with 10 % v/v fetal bovine serum (FBS) and 1 % v/v Pen/Strep (DMEM, VWR, Radnor, PA, USA), following standard practices. All chemicals, mediums, and substrates were purchased from VWR (Radnor, PA, USA), otherwise mentioned.

### Material Preparation

GelMA was synthesized according to an established protocol.^49^ In a 100 mL glass flask, Dulbecco’s phosphate-buffered saline (DPBS) (Sigma-Aldrich, St. Louis, MO, USA) was mixed with 10% w/v porcine skin gelatin (CAS Number 9000–70–8; Sigma-Aldrich). The flask was covered to prevent evaporation and stirred using a magnetic stir bar on a hot plate at 60 °C until fully dissolved (approximately one hour). After the gelatin was completely dissolved in the DPBS, 5 mL of methacrylic anhydride (CAS: 760–93–0, Sigma-Aldrich) was slowly pipetted into the solution. The temperature was then lowered to 50 °C, and the solution was stirred and allowed to react for one hour. To stop the reaction, pre-warmed DPBS—five times the volume of the initial solution—was added. GelMA was dialyzed using dialysis tubing (12–14 kDa molecular weight cut-off) for one week to remove excess methacrylic anhydride. The resulting solution was transferred into centrifuge tubes and stored at −80 °C.

Hyaluronic acid (BioSynthesis Inc., Germany) was dissolved in PBS (2 g in 100 mL) and stirred at 4 °C until completely dissolved. Then, 67 mL of N,N-dimethylformamide (DMF, Sigma) was added to the hyaluronic acid solution and stirred until well mixed. An appropriate amount of methacrylic anhydride (CAS: 760–93–0, Sigma-Aldrich, St. Louis, MO, USA) was added dropwise, and 1 M sodium hydroxide (NaOH, Sigma) was used to adjust the pH. The solution was reacted at 4 °C for 24 hours, then mixed with 95% ethanol to precipitate and crystallize the product. The precipitate was washed with ethanol and deionized water and subsequently dissolved in deionized water. The solution was dialyzed using 12–14 kDa dialysis tubing for one week. The NMR spectrum was then used to confirm the efficiency of chemical yield and the degree of acrylation. After freeze-drying, a HAMA stock solution (2% w/v) was prepared in DPBS and pre-warmed to 50 °C with constant stirring. Once fully dissolved, solutions of 7% GelMA and 3% GelMA with 1% HAMA were prepared from the stock solutions. After continued stirring at 50 °C, lithium phenyl-2,4,6-trimethylbenzoylphosphinate (LAP, Sigma) at 1% w/v was added as the photoinitiator. All solutions were filtered using syringe filters.

### Bioink Preparation and Biofabrication

GelMA and HAMA/GelMA solutions were mixed with pelleted cells at the desired concentration (1 million cells/mL) to form cellular hydrogels. Acellular hydrogels were prepared in the same way, without the addition of cells. Using our extrusion bioprinter, we printed predefined models and then crosslinked the constructs using post-extrusion light exposure.^26^ Disk-shaped samples (8 mm in diameter and 2.5 mm in height) were fabricated in triplicate for each time point (Days 0, 1, 3, 7, 14, and 21). Samples were crosslinked under UV light for 3 minutes at an intensity of 3800 × 100 μJ/cm^2^ (VWR).

After crosslinking, cellular samples were incubated in DMEM supplemented with 10% v/v fetal bovine serum (FBS) and 1% v/v Penicillin-Streptomycin (Pen/Strep) (VWR, Radnor, PA, USA), while acellular samples were incubated in DPBS. All samples were maintained in anti-adherent coated (Anti-Adherence Rinsing Solution, AggreWell)12-well plates.

### Mechanical Characterization

Standard compression testing was performed to measure the stiffness under compression load. Samples were placed on a metal flat plate of the universal testing machine (Instron, MA, USA), and the tests were performed using a strain rate of 0.5 mm/min. Regression was used to determine the linear area, and Young’s modulus was calculated as the slope of the stress-strain curve from 0-10 % strain.

### Mass Characterization

After mechanical characterization was completed, samples were massed with a balance scale. Samples were fixed in 4% paraformaldehyde (Thermo Fisher) for 2 h. Paraformaldehyde was removed, and the samples were kept at 4 °C in DPBS. Samples were frozen at −80°C for 24 h and lyophilized for 48 h. Mass residuals were calculated using the mass of the chosen time point (M_x_) divided by the average mass of day 0 samples (M_0_):

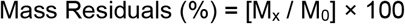

### Swelling Measurement

Fixed samples were weighed again and placed in centrifuge tubes with perforated caps to allow airflow. The samples were then freeze-dried to produce dry scaffolds. These dry scaffolds were weighed using a balance scale. The swelling ratio was calculated using the following formula, where M_w_ is the mass of the wet sample and M_d_ is the mass of the corresponding dry sample:^26,50^

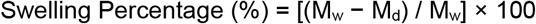

### Porosity and Pore Size Measurement

Scaffolds (*n* = 5) were measured with a digital micrometer (Mitutoyo, Japan), and the volume was calculated. The dry scaffolds were weighed with an electronic analytical balance (ML54T, Mettler Toledo, Switzerland) and immersed in 30 ml of isopropanol with a density of 0.785 g/ml under vacuum overnight to remove the air in the pores. The wet scaffold was weighed, and the porosity was calculated. The samples were freeze-dried, attached to the Scanning Electron Microscopy (SEM) stubs, and coated with gold for 30 s under a vacuum. Samples were imaged using Zeiss Supra V 40VP EM under a high vacuum. Images were analyzed using ImageJ for pore size measurements.

### Static Diffusion

Fixed hydrogels were rewashed with DI water three times for 20 minutes each, then cut into smaller pieces and weighed. The hydrogels were immersed in Rhodamine B (RhB) at a concentration of 0.5 mg/mL for 3 minutes, after which they were removed, washed twice with DI water to eliminate surface-bound RhB, and placed into a 24-well plate or separate beakers containing 1 mL of DI water overnight. The following day, the samples were measured at a fluorescent excitation peak at 546 nm and an emission peak at 567 nm. The gel samples were then immersed in a RhB bath (0.5 mg/mL) for 2 hours to reach a steady state, and the release measurement step was repeated. We assumed 1-D diffusion transport and used Fick’s law to calculate the diffusion coefficient.^46^

### Infrared Analysis

Fourier Transform Infrared-Attenuated Total Reflection (FTIR-ATR) spectra of GelMA and HAMA/GelMA were obtained on a Perkin Elmer FT-IR System Spectrum BX spectrophotometer (Perkin-Elmer Inc., USA) equipped with a single horizontal GoldenGate ATR cell. To obtain each spectrum, 64 scans were acquired in the 4000–500 cm^−1^ range, with a resolution of 4 cm^−1^.

### Cell Viability and Metabolic Activity

Cellular responses to GelMA and HAMA/GelMA hydrogels were evaluated using Live/Dead staining (Invitrogen™ LIVE/DEAD™ Viability/Cytotoxicity Kit) and the CCK8 assay (Sigma-Aldrich). A Zeiss Axio Imager Z1 microscope was used for imaging. ImageJ software was employed to quantify live and dead cells and calculate cell viability as a percentage. The CCK8 assay was used to assess the metabolic activity of encapsulated cells. Results were normalized to negative controls and expressed as optical density (OD) values.

### Immunostaining

The samples were fixed in 4% (v/v) paraformaldehyde and washed three times with 1× PBS. They were then permeabilized using 0.1% Triton X-100 (Sigma-Aldrich) in DPBS for 15 minutes, followed by blocking with 1% bovine serum albumin (BSA) blocking buffer (Alfa Aesar, Haverhill, MA) for 1 hour. Samples were subsequently incubated overnight at 4°C with 1:100 FITC-conjugated phalloidin (1 μg/mL, Millipore Sigma, FAK100) in 1% BSA. Simultaneous permeabilization and blocking were also performed prior to incubation with the primary antibody (anti-Ki67; 1:500, Abcam, USA), which was carried out overnight at 4°C. Hydrogels were counterstained with DAPI (0.02 mg/mL, ThermoFisher Scientific, USA), mounted, and cover-slipped prior to imaging with a Zeiss Axio Imager.Z1 microscope. The percentage of proliferating cells (Ki67-positive) was calculated as follows:

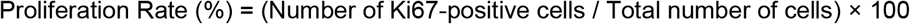

Hydrogels were subdivided into center and edge regions. The edge region was defined as the area within 0.5 mm of the hydrogel perimeter, while the center region included the area within a 2 mm radius from the geometric center of the cylindrical hydrogel. Quantification of immunofluorescent staining intensity was performed using the ImageJ software analysis tool. The mean staining intensity (pixel density) was normalized to the number of cells within each field of view. In addition, spheroid size, distribution, and roundness were assessed from three representative images per group and time point.

### Theoretical Model

A theoretical model was developed to estimate the total energy of the cell and the ECM. This model was implemented within a finite element framework. Using this framework, we simulated each of the three possible cell morphological states—rounded, elongated, and clustered—within both soft and stiff ECMs, resulting in a total of six cases. For each case, the total energy was calculated. Detailed descriptions of the model and finite element simulations are provided in Supplementary Notes 1-3. The code implementing the model in the finite element framework is available on GitHub (https://github.com/Farid-Alisafaei/ECM-Mechanics-Regulate-Cancer-Cell-State) and includes annotations describing the model parameters.

### Statistical Analysis

Statistical analysis was performed using Microsoft Excel and the GraphPad Prism9 statistical tool. A one-way analysis of variance (one-way ANOVA and two-way ANOVA) test was used for data analysis. The value of *p* < 0.05 was considered to be statistically significant.

## Acknowledgment

The authors gratefully acknowledge NJIT funding, and A.K.M. acknowledges the financial support from NSF-2243506 and NSF-2426919 grants for this work.

## Conflict of Interest

The authors declare no conflict of interest.

## Notes

### Competing Interest Statement

The authors have declared no competing interest.

