## Supplementary Information for "Extracellular Matrix Physical Properties Regulate Cancer Cell Morphological Transitions in 3D Hydrogel Microtissues"

### 1. Three-dimensional cytoskeletal model

In this section, we describe the theoretical cytoskeleton model used in our simulations. We developed a 3D continuum-based constitutive model in which the total cytoskeletal response arises from the coupled contributions of (i) myosin molecular motors, (ii) actin filaments, and (iii) microtubules, as illustrated schematically in Fig. 2B. This constitutive model was implemented into a finite element framework, such that each element follows the same underlying material law derived from these cytoskeletal components. Using this implementation, we simulated the three possible cell morphological states and computed the associated energy components, as described in Supplementary Information (SI) Sections 2 and 3.

#### 1.1 Myosin motors, actin filaments, and microtubules

The first component of the cytoskeletal model is myosin, which generates cell internal contractility and is denoted by a contractility tensor, $\rho_{ij}$, whose components represent cell internally-generated stresses in different directions.[1] In what follows, we first define the contractility tensor $\rho_{ij}$ and we then discuss how the cell model responds to mechanical signals from their extracellular environment by adjusting $\rho_{ij}$.

The force generated by a myosin motor at submicron levels can be treated as a force dipole Fig. 2B. Force dipoles are pairs of equal but oppositely directed forces $F_i(x_j)$ and $-F_i(x_j + \Delta x_j)$ where $|\Delta x_j|$ is the length of myosin II filaments (approximately 200 nm),[2] and $x_j$ and $x_j + \Delta x_j$ are coordinates of myosin head domains. We then can define the work done by a force dipole $W_{\text{dipole}}$ and the total work generated by all myosin motors per volume $W$ as follows, respectively,

$$W_{\text{dipole}} = F_i u_i(x_j + \Delta x_j) - F_i u_i(x_j) \tag{S1.1}$$

$$W = \left(\frac{1}{V}\right) \sum_{k=1}^{N} \left[F_i^{(k)} u_i\left(x_j^{(k)} + \Delta x_j^{(k)}\right) - F_i^{(k)} u_i\left(x_j^{(k)}\right)\right] \tag{S1.2}$$

where $u_i(x_j)$ and $u_i(x_j + \Delta x_j)$ are the displacements of the cytoskeleton at $x_j$ and $x_j + \Delta x_j$, respectively, and $V$ is the volume of the cytoplasm. Assuming that $N$ is the total number of bound (phosphorylated) myosin motors and using equation (S1.4), we can rewrite equation (S1.2) in this form

$$W = \left(\frac{1}{V}\right) \sum_{k=1}^{N} F_i^{(k)} \Delta x_j^{(k)} \partial_j u_i \tag{S1.3}$$

$$\partial_j u_i = \frac{u_i\left(x_j^{(k)} + \Delta x_j^{(k)}\right) - u_i\left(x_j^{(k)}\right)}{\Delta x_j^{(k)}} \tag{S1.4}$$

Therefore, the total work can be presented as a function of the contractility tensor $\rho_{ij}$

$$W = \rho_{ij} \varepsilon_{ij} \tag{S1.5}$$

where

$$\varepsilon_{ij} = \frac{1}{2}\left(\partial_j u_i + \partial_i u_j\right) \tag{S1.6}$$

is the linearized strain tensor, and

$$\rho_{ij} = \left(\frac{1}{V}\right)\sum_{k=1}^{N} F_i^{(k)}\,\Delta x_j^{(k)} \tag{S1.7}$$

Note that the contractility tensor $\rho_{ij}$ is a symmetric tensor, satisfying the following condition,

$$F_i^{(k)}\Delta x_j^{(k)} = F_j^{(k)}\Delta x_i^{(k)} \tag{S1.8}$$

Equation (S1.7) demonstrates that the cell contractility tensor $\rho_{ij}$ represents the density of phosphorylated myosin molecular motors. Experimental observations have shown that cells adjust the density of phosphorylated myosin in response to mechanical signals from the environment. For example, in 2D, cells on stiff substrates have higher densities of phosphorylated myosin and are more contractile than those on soft substrates.[3,4] Therefore, to have a contractile model that actively responds to extracellular mechanical signals, we hypothesize that the average of cell contractility in all three directions, $\frac{1}{3}\rho_{kk} = (\rho_{11} + \rho_{22} + \rho_{33})/3$, in our coarse-grain model increases with the average of tension in the actin filament network, $\frac{1}{3}\sigma_{kk} = (\sigma_{11} + \sigma_{22} + \sigma_{33})/3$

$$\frac{\rho_{kk}}{3} = f_{\mathrm{m}}\frac{\sigma_{kk}}{3} + f_0\,\rho_0 \tag{S1.9}$$

where this tension-dependent feedback mechanism is regulated by the feedback constant parameter $f_{\mathrm{m}}$. In equation (S1.9), the constant parameter $\rho_0$ is the initial cell contractility, and the constant parameter $f_0$ regulates the mean contractility $\frac{1}{3}\rho_{kk}$ in the absence of tension ($\sigma_{kk} = 0$). We have previously shown that as a result of the feedback mechanism in (S1.9), in addition to the cell contractility $\rho_{ij}$, the stiffness of the actin network $C_{ijkl}^{(A)}$ and the tension it carries, $\sigma_{ij}$, also increase with substrate stiffness in an orientation-dependent manner.[5] We will later show how the feedback between $\rho_{kk}$ and $\sigma_{kk}$, introduced in Eq. S1.9, emerges naturally from our model, as made explicit in the expanded form presented in Eq. S1.54.

To be able to implement the model in (S1.9) into a three-dimensional finite element model, we need to derive the total stress $\sigma_{ij}$ and total stiffness $C_{ij}$ tensors. To this end, we start with the following definition for $\rho_{ij}$ which relates it to the strain tensor $\boldsymbol{\varepsilon}^{(X)}$ (the three-dimensional representation of $\varepsilon^{(X)}$ shown in Fig. S7)

$$\rho_{ij} = K^{(\rho)}\varepsilon_{kk}^{(X)}\delta_{ij} + 2\mu^{(\rho)}\left(\varepsilon_{ij}^{(X)} - \tfrac{1}{3}\varepsilon_{kk}^{(X)}\delta_{ij}\right) + \bar{\rho}_0\delta_{ij} \tag{S1.10}$$

where

$$K^{(\rho)} = \frac{3K^{(\mathrm{MT})}\alpha_{\mathrm{v}} - 1}{3(\beta_{\mathrm{v}} - \alpha_{\mathrm{v}})} \tag{S1.11}$$

is effective modulus for the motor density,

$$\mu^{(\rho)} = \frac{2\mu^{(\mathrm{MT})}\alpha_{\mathrm{d}} - 1}{2(\beta_{\mathrm{d}} - \alpha_{\mathrm{d}})} \tag{S1.12}$$

is the effective modulus of the polarization,

$$\bar{\rho}_0 = \frac{\beta_{\mathrm{v}}\,\rho_0}{\beta_{\mathrm{v}} - \alpha_{\mathrm{v}}} \tag{S1.13}$$

is the effective contractility,

$$K^{(\mathrm{MT})} = \frac{E^{(\mathrm{MT})}}{3(1 - 2v^{(\mathrm{MT})})} \tag{S1.14}$$

is the bulk modulus of the cytoskeletal components that are in compression (*e.g.*, the microtubule network), and

$$\mu^{(\mathrm{MT})} = \frac{E^{(\mathrm{MT})}}{2(1 + v^{(\mathrm{MT})})} \tag{S1.15}$$

is the shear modulus of the cytoskeletal components that are in compression (*e.g.*, the microtubule network). $E^{(\mathrm{MT})}$ in these equations is the elastic modulus of the cytoskeletal components that are in compression (*e.g.*, the microtubule network), $v^{(\mathrm{MT})}$ is the Poisson's ratio of the cytoskeletal components that are in compression (*e.g.*, the microtubule network), $\rho_0$ is the baseline cell contractility, $\alpha_{\mathrm{v}}$ is the volumetric chemo-mechanical feedback parameter (large values of $\alpha_{\mathrm{v}}$ lead to higher phosphorylation of myosin), $\alpha_{\mathrm{d}}$ is the deviatoric chemo-mechanical feedback parameter which represents the cell's tendency to polarized contraction (small values of $\alpha_{\mathrm{d}}$ results in non-polarized contractility), $\beta_{\mathrm{v}}$ is the volumetric chemical stiffness parameter which regulates the mean contractility $\frac{1}{3}\rho_{kk}$ in the absence of tension ($\sigma_{kk} = 0$) by maintaining the cell contractility at its initial level (large values of $\beta_{\mathrm{v}}$ keeps the level of myosin phosphorylation low), and $\beta_{\mathrm{d}}$ is the deviatoric chemical stiffness parameter which represents the disinclination of myosin to orient along the cell's polarization direction (large values of $\beta_{\mathrm{d}}$ leads to a random orientation of myosin motors).

A large portion of microtubules experiences compression due to the cell's internal contractile forces.[6–8] Consistently, the cell contractility $\rho_{ij}$ in our model generates the compressive stress $C_{ijkl}^{(\mathrm{MT})}\varepsilon_{kl}^{(\mathrm{X})}$ on microtubules where

$$C_{ijkl}^{(\mathrm{MT})} = K^{(\mathrm{MT})}\delta_{ij}\delta_{kl} + \mu^{(\mathrm{MT})}\left(\delta_{ik}\delta_{jl} + \delta_{il}\delta_{jk} - \frac{2}{3}\delta_{ij}\delta_{kl}\right) \tag{S1.16}$$

is the microtubule network stiffness tensor.

Unlike microtubules, actin filaments experience tension. Similarly, the cell contractility $\rho_{ij}$ in our model generates tension in the actin network, $\sigma_{ij}$, in addition to compressing microtubules (Fig. S7),

$$\rho_{ij} = -C^{(\mathrm{MT})}_{ijkl}\varepsilon^{(\mathrm{X})}_{kl} + \sigma_{ij} \tag{S1.17}$$

In our model, Equation (S1.10) indicates that the contractility tensor $\rho_{ij}$ is initially isotropic and, as a result, the cell exhibits the same contractility in all directions in the initial configuration. This can be mathematically shown by rewriting equation (S1.10) in the following form

$$\rho_{ij} = C^{(\rho)}_{ijkl}\varepsilon^{(\mathrm{X})}_{kl} + \bar{\rho}_0\delta_{ij} \tag{S1.18}$$

where

$$C^{(\rho)}_{ijkl} = K^{(\rho)}\delta_{ij}\delta_{kl} + \mu^{(\rho)}\left(\delta_{ik}\delta_{jl} + \delta_{il}\delta_{jk} - \frac{2}{3}\delta_{ij}\delta_{kl}\right) \tag{S1.19}$$

However, the model predicts that the contractility can become nonuniform and anisotropic, dependent on extracellular physical constraints.[5,9] For all simulations, we start with a uniform (independent of spatial location) and isotropic (independent of direction) contractility. Equations (S1.17) and (S1.18) demonstrate that, in the stress-free condition $\sigma_{ij} = 0$ (initially there is no tension), the diagonal components of $\rho_{ij}$ are all equal and non-zero $\rho_{11} = \rho_{22} = \rho_{33} \neq 0$, while the off-diagonal components are all zero $\rho_{12} = \rho_{21} = \rho_{13} = \rho_{31} = \rho_{23} = \rho_{32} = 0$, indicating that the contractility tensor $\rho_{ij}$ is initially isotropic. This is better shown by writing equation (S1.18) in the following format using Voigt notation

$$\begin{Bmatrix}\rho_{11}\\ \rho_{22}\\ \rho_{33}\\ \rho_{12}\\ \rho_{13}\\ \rho_{23}\end{Bmatrix} = \begin{bmatrix} C^{(\rho)}_{1111} & C^{(\rho)}_{1122} & C^{(\rho)}_{1133} & C^{(\rho)}_{1112} & C^{(\rho)}_{1113} & C^{(\rho)}_{1123}\\ C^{(\rho)}_{2211} & C^{(\rho)}_{2222} & C^{(\rho)}_{2233} & C^{(\rho)}_{2212} & C^{(\rho)}_{2213} & C^{(\rho)}_{2223}\\ C^{(\rho)}_{3311} & C^{(\rho)}_{3322} & C^{(\rho)}_{3333} & C^{(\rho)}_{3312} & C^{(\rho)}_{3313} & C^{(\rho)}_{3323}\\ C^{(\rho)}_{1211} & C^{(\rho)}_{1222} & C^{(\rho)}_{1233} & C^{(\rho)}_{1212} & C^{(\rho)}_{1213} & C^{(\rho)}_{1223}\\ C^{(\rho)}_{1311} & C^{(\rho)}_{1322} & C^{(\rho)}_{1333} & C^{(\rho)}_{1312} & C^{(\rho)}_{1313} & C^{(\rho)}_{1323}\\ C^{(\rho)}_{2311} & C^{(\rho)}_{2322} & C^{(\rho)}_{2333} & C^{(\rho)}_{2312} & C^{(\rho)}_{2313} & C^{(\rho)}_{2323}\end{bmatrix} \begin{Bmatrix}\varepsilon_{11}\\ \varepsilon_{22}\\ \varepsilon_{33}\\ \varepsilon_{12}\\ \varepsilon_{13}\\ \varepsilon_{23}\end{Bmatrix} + \begin{Bmatrix}\bar{\rho}_0\\ \bar{\rho}_0\\ \bar{\rho}_0\\ 0\\ 0\\ 0\end{Bmatrix} \tag{S1.20}$$

where the first, second, and third components of the last term in equation (S1.20) have the same value $\bar{\rho}_0$.

Concomitant with the phosphorylation of more myosin, cells also respond to tension by polymerizing actin filaments and bundling and aligning them in the direction of tension.[10,11] Thus, in addition to the feedback mechanism between the contractility $\rho_{ij}$ and the stress $\sigma_{ij}$ in (S1.9), we also hypothesize that the stiffness of the actin network $C^{(\mathrm{A})}_{ijkl}$ increases in proportion to, and in the directions of, the tensile principal components of the stress tensor $\sigma_{ij}$,[5]

$$\mathrm{C}_{ijkl}^{(\mathrm{A})} = \mathrm{C}_{ijkl}^{(\mathrm{I})} + \mathrm{C}_{ijkl}^{(\mathrm{F})} \tag{S1.21}$$

In Equation (S1.21), $\mathbf{C}^{(\mathrm{F})}$ denotes the stiffening of the actin network with tension (but not in compression) and $\mathbf{C}^{(\mathrm{I})}$ is the initial stiffness of the actin filaments network

$$\mathrm{C}_{ijkl}^{(\mathrm{I})} = K^{(\mathrm{I})}\delta_{ij}\delta_{kl} + \mu^{(\mathrm{I})}\left(\delta_{ik}\delta_{jk} + \delta_{il}\delta_{jk} - \frac{2}{3}\delta_{ij}\delta_{kl}\right) \tag{S1.22}$$

where

$$K^{(\mathrm{I})} = \frac{E^{(\mathrm{I})}}{3(1 - 2v^{(\mathrm{I})})} \tag{S1.23}$$

is the initial bulk modulus of the actin network, and

$$\mu^{(\mathrm{I})} = \frac{E^{(\mathrm{I})}}{2(1 + v^{(\mathrm{I})})} \tag{S1.24}$$

is the initial shear modulus of the actin network, $E^{(\mathrm{I})}$ is the initial elastic modulus of the actin network, and $v^{(\mathrm{I})}$ is the initial Poisson's ratio of the actin network.

To define the stiffening part of the stiffness tensor, $\mathbf{C}^{(\mathrm{F})}$, we first decompose $\boldsymbol{\sigma}$

$$\sigma_{ij} = \sigma_{ij}^{(\mathrm{A})} = \sigma_{ij}^{(\mathrm{I})} + \sigma_{ij}^{(\mathrm{F})} \tag{S1.25}$$

where $\boldsymbol{\sigma}^{(\mathrm{I})}$

$$\sigma_{ij}^{(\mathrm{I})} = \mathrm{C}_{ijkl}^{(\mathrm{I})}\,\varepsilon_{kl}^{(\mathrm{Y})} \tag{S1.26}$$

is linearly related to the strain tensor $\boldsymbol{\varepsilon}^{(\mathrm{Y})}$ (the three-dimensional representation of $\varepsilon^{(\mathrm{Y})}$ shown in Supplementary Figure S7) which can be written as a function of its eigenvalues (principal strains) $\varepsilon_1^{(\mathrm{Y})}$, $\varepsilon_2^{(\mathrm{Y})}$, $\varepsilon_3^{(\mathrm{Y})}$ and eigenvectors $\boldsymbol{n}_1$, $\boldsymbol{n}_2$, $\boldsymbol{n}_3$

$$\boldsymbol{\varepsilon}^{(\mathrm{Y})} = \sum_{i=1}^{3} \varepsilon_i^{(\mathrm{Y})}\,\boldsymbol{n}_i \otimes \boldsymbol{n}_i = \sum_{i=1}^{3} \varepsilon_i^{(\mathrm{Y})}\,\mathbf{E}_i \tag{S1.27}$$

where the symmetric tensors $\mathbf{E}_1 = \boldsymbol{n}_1 \otimes \boldsymbol{n}_1$, $\mathbf{E}_2 = \boldsymbol{n}_2 \otimes \boldsymbol{n}_2$, and $\mathbf{E}_3 = \boldsymbol{n}_3 \otimes \boldsymbol{n}_3$ are the eigenprojections of $\boldsymbol{\varepsilon}^{(\mathrm{Y})}$ and $\otimes$ denotes the dyadic product of two arbitrary vectors $\boldsymbol{u}$ and $\boldsymbol{v}$ as $(\boldsymbol{u} \otimes \boldsymbol{v})_{ij} = u_i v_j$. With the eigenvalues ($\varepsilon_1^{(\mathrm{Y})}$, $\varepsilon_2^{(\mathrm{Y})}$, $\varepsilon_3^{(\mathrm{Y})}$) and eigenvectors ($\boldsymbol{n}_1$, $\boldsymbol{n}_2$, $\boldsymbol{n}_3$) at hand, we next define $\sigma_{ij}^{(\mathrm{F})}$ in equation (S1.25)

$$\boldsymbol{\sigma}^{(\mathrm{F})} = \sum_{i=1}^{3} \frac{\partial f(\varepsilon_i^{(\mathrm{Y})})}{\partial \varepsilon_i^{(\mathrm{Y})}} \boldsymbol{n}_i \otimes \boldsymbol{n}_i = \sum_{i=1}^{3} \sigma^{(\mathrm{F})}\left(\varepsilon_i^{(\mathrm{Y})}\right)\mathbf{E}_i = \sum_{i=1}^{3} \sigma_i^{(\mathrm{F})}\mathbf{E}_i \tag{S1.28}$$

where $\sigma_i^{(\mathrm{F})}$ are the eigenvalues (principal stresses) of the stress tensor $\sigma_{ij}^{(\mathrm{F})}$ and are defined as follows

$$\sigma_i^{(\mathrm{F})} = \frac{\partial f(\varepsilon_i^{(\mathrm{Y})})}{\partial \varepsilon_i^{(\mathrm{Y})}} = \tag{S1.29}$$

$$\begin{cases} 0 & \varepsilon_i^{(\mathrm{Y})} < \epsilon_1 \\ \ell \dfrac{\left(\dfrac{\varepsilon_i^{(\mathrm{Y})} - \epsilon_1}{\epsilon_2 - \epsilon_1}\right)^t \left(\varepsilon_i^{(\mathrm{Y})} - \epsilon_1\right)^2}{(t+1)(t+2)} & \epsilon_1 \le \varepsilon_i^{(\mathrm{Y})} < \epsilon_2 \\ \ell \left[\dfrac{\left(1 + \varepsilon_i^{(\mathrm{Y})} - \epsilon_2\right)^{s+2} - 1}{(s+1)(s+2)} + \dfrac{\epsilon_2 - \varepsilon_i^{(\mathrm{Y})}}{s+1} + \dfrac{(\varepsilon_i^{(\mathrm{Y})} - \epsilon_2)(\epsilon_2 - \epsilon_1)}{t+1} + \dfrac{(\epsilon_2 - \epsilon_1)^2}{(t+1)(t+2)}\right] & \varepsilon_i^{(\mathrm{Y})} \ge \epsilon_2 \end{cases}$$

to ensure the continuity and smoothness of the first and second derivatives of $\sigma_i^{(\mathrm{F})}$ with respect to $\varepsilon_i^{(\mathrm{Y})}$ at the transition points $\epsilon_1 = \epsilon_\mathrm{c} - 0.5\epsilon_\mathrm{t}$ and $\epsilon_2 = \epsilon_\mathrm{c} + 0.5\epsilon_\mathrm{t}$ where $\epsilon_\mathrm{t} = 0.25\epsilon_\mathrm{c}$ is the transition width, $\epsilon_\mathrm{c}$ is the critical (tensile) principal strain, and $t$ is the transition constant. Equation (S1.29) shows that for large tensile strains $\varepsilon_i^{(\mathrm{Y})} \ge \epsilon_2$, the principal stress $\sigma_i^{(\mathrm{F})}$ nonlinearly increases with the principal strain $\varepsilon_i^{(\mathrm{Y})}$ where this increase is regulated by the stiffening parameters $\ell$ and $s$. With $\boldsymbol{\sigma}^{(\mathrm{F})}$ at hand from equation (S1.28), we can now determine $\mathbf{C}^{(\mathrm{F})}$ as follows

$$\mathrm{C}_{ijkl}^{(\mathrm{F})} = \frac{\mathrm{d}\sigma_{ij}^{(\mathrm{F})}}{\mathrm{d}\varepsilon_{kl}^{(\mathrm{Y})}} \qquad \text{or} \qquad \mathbf{C}^{(\mathrm{F})} = \frac{\mathrm{d}\boldsymbol{\sigma}^{(\mathrm{F})}}{\mathrm{d}\boldsymbol{\varepsilon}^{(\mathrm{Y})}} \tag{S1.30}$$

The piecewise linear approximation in equation (S1.30) requires $\sigma_{ij}^{(\mathrm{F})}$ as a function of $\varepsilon_{ij}^{(\mathrm{Y})}$ while equation (S1.29) gives $\sigma_i^{(\mathrm{F})}$ as a function of $\varepsilon_i^{(\mathrm{Y})}$. Thus, we first write $\mathrm{C}_{ijkl}^{(\mathrm{F})}$ in the following form using the definition of $\sigma_{ij}^{(\mathrm{F})}$ in equation (S1.28)

$$\mathbf{C}^{(\mathrm{F})} = \sum_{i=1}^{3} \left\{ \mathbf{E}_i \otimes \frac{\mathrm{d}\sigma_i^{(\mathrm{F})}}{\mathrm{d}\boldsymbol{\varepsilon}^{(\mathrm{Y})}} + \sigma_i^{(\mathrm{F})} \frac{\mathrm{d}\mathbf{E}_i}{\mathrm{d}\boldsymbol{\varepsilon}^{(\mathrm{Y})}} \right\} \tag{S1.31}$$

and we then expand the first term in the right-hand side of equation (S1.31) by applying the chain rule

$$\mathbf{C}^{(\mathrm{F})}=\sum_{i=1}^{3}\left\{\sum_{j=1}^{3}\frac{\partial\sigma_i^{(\mathrm{F})}}{\partial\varepsilon_j^{(\mathrm{Y})}}\mathbf{E}_i\otimes\frac{\mathrm{d}\varepsilon_j^{(\mathrm{Y})}}{\mathrm{d}\boldsymbol{\varepsilon}^{(\mathrm{Y})}}+\sigma_i^{(\mathrm{F})}\frac{\mathrm{d}\mathbf{E}_i}{\mathrm{d}\boldsymbol{\varepsilon}^{(\mathrm{Y})}}\right\} \tag{S1.32}$$

To further expand equation (1.32) and derive an exact form for $\mathbf{C}^{(\mathrm{F})}$, we consider the three following cases.

In the first case, the three eigenvalues of the strain tensor $\varepsilon_{ij}^{(\mathrm{Y})}$ are all nonidentical ($\varepsilon_1^{(\mathrm{Y})}\neq\varepsilon_2^{(\mathrm{Y})}\neq\varepsilon_3^{(\mathrm{Y})}$). In this case, we can derive the following expression for $\mathbf{C}^{(\mathrm{F})}$ from equation (S1.32) by taking the derivatives of $\varepsilon_j^{(\mathrm{Y})}$ and $\mathbf{E}_i$ with respect to $\boldsymbol{\varepsilon}^{(\mathrm{Y})}$

$$\begin{aligned}\mathbf{C}^{(\mathrm{F})}=\sum_{a=1}^{3}\frac{\sigma_a^{(\mathrm{F})}}{(\varepsilon_a^{(\mathrm{Y})}-\varepsilon_b^{(\mathrm{Y})})(\varepsilon_a^{(\mathrm{Y})}-\varepsilon_c^{(\mathrm{Y})})}&\left\{\frac{\mathrm{d}\left(\boldsymbol{\varepsilon}^{(\mathrm{Y})}\right)^2}{\mathrm{d}\boldsymbol{\varepsilon}^{(\mathrm{Y})}}-\left(\varepsilon_b^{(\mathrm{Y})}+\varepsilon_c^{(\mathrm{Y})}\right)\mathbf{I}_\mathrm{S}\right.\\&-\left[\left(\varepsilon_a^{(\mathrm{Y})}-\varepsilon_b^{(\mathrm{Y})}\right)+\left(\varepsilon_a^{(\mathrm{Y})}-\varepsilon_c^{(\mathrm{Y})}\right)\right]\mathbf{E}_a\otimes\mathbf{E}_a\\&\left.-\left(\varepsilon_b^{(\mathrm{Y})}-\varepsilon_c^{(\mathrm{Y})}\right)(\mathbf{E}_b\otimes\mathbf{E}_b-\mathbf{E}_c\otimes\mathbf{E}_c)\right\}+\sum_{i=1}^{3}\sum_{j=1}^{3}\frac{\partial\sigma_i^{(\mathrm{F})}}{\partial\varepsilon_j^{(\mathrm{Y})}}\mathbf{E}_i\otimes\mathbf{E}_j\end{aligned} \tag{S1.33}$$

where

$$\left(\frac{\mathrm{d}\left(\boldsymbol{\varepsilon}^{(\mathrm{Y})}\right)^2}{\mathrm{d}\boldsymbol{\varepsilon}^{(\mathrm{Y})}}\right)_{ijkl}=\frac{1}{2}\left(\delta_{ik}\varepsilon_{lj}^{(\mathrm{Y})}+\delta_{il}\varepsilon_{kj}^{(\mathrm{Y})}+\delta_{jl}\varepsilon_{ik}^{(\mathrm{Y})}+\delta_{kj}\varepsilon_{il}^{(\mathrm{Y})}\right) \tag{S1.34}$$

is the derivative of the square of $\boldsymbol{\varepsilon}^{(\mathrm{Y})}$, and

$$(\mathbf{I}_\mathrm{S})_{ijkl}=\frac{1}{2}\left(\delta_{ik}\delta_{jl}+\delta_{il}\delta_{jk}\right) \tag{S1.35}$$

is the symmetric identity tensor.

In the second case, $\varepsilon_{ij}^{(\mathrm{Y})}$ has two identical eigenvalues ($\varepsilon_1^{(\mathrm{Y})}\neq\varepsilon_2^{(\mathrm{Y})}=\varepsilon_3^{(\mathrm{Y})}$) which gives the following analytical expression for $\mathbf{C}^{(\mathrm{F})}$

$$\mathbf{C}^{(\mathrm{F})}=s_1\frac{\mathrm{d}\left(\boldsymbol{\varepsilon}^{(\mathrm{Y})}\right)^2}{\mathrm{d}\boldsymbol{\varepsilon}^{(\mathrm{Y})}}-s_2\mathbf{I}_\mathrm{S}-s_3\boldsymbol{\varepsilon}^{(\mathrm{Y})}\otimes\boldsymbol{\varepsilon}^{(\mathrm{Y})}+s_4\boldsymbol{\varepsilon}^{(\mathrm{Y})}\otimes\mathbf{I}+s_5\mathbf{I}\otimes\boldsymbol{\varepsilon}^{(\mathrm{Y})}-s_6\mathbf{I}\otimes\mathbf{I} \tag{S1.36}$$

where

$$\mathrm{I}_{ij}=\delta_{ij} \tag{S1.37}$$

is the second-order identity tensor, and

$$s_1 = \frac{\sigma_a^{(\mathrm{F})} - \sigma_c^{(\mathrm{F})}}{\left(\varepsilon_a^{(\mathrm{Y})} - \varepsilon_c^{(\mathrm{Y})}\right)^2} + \frac{1}{\varepsilon_a^{(\mathrm{Y})} - \varepsilon_c^{(\mathrm{Y})}}\left(\frac{\partial \sigma_c^{(\mathrm{F})}}{\partial \varepsilon_b^{(\mathrm{Y})}} - \frac{\partial \sigma_c^{(\mathrm{F})}}{\partial \varepsilon_c^{(\mathrm{Y})}}\right) \tag{S1.38a}$$

$$s_2 = 2\varepsilon_c^{(\mathrm{Y})} \frac{\sigma_a^{(\mathrm{F})} - \sigma_c^{(\mathrm{F})}}{\left(\varepsilon_a^{(\mathrm{Y})} - \varepsilon_c^{(\mathrm{Y})}\right)^2} + \frac{\varepsilon_a^{(\mathrm{Y})} + \varepsilon_c^{(\mathrm{Y})}}{\varepsilon_a^{(\mathrm{Y})} - \varepsilon_c^{(\mathrm{Y})}}\left(\frac{\partial \sigma_c^{(\mathrm{F})}}{\partial \varepsilon_b^{(\mathrm{Y})}} - \frac{\partial \sigma_c^{(\mathrm{F})}}{\partial \varepsilon_c^{(\mathrm{Y})}}\right) \tag{S1.38b}$$

$$s_3 = 2\frac{\sigma_a^{(\mathrm{F})} - \sigma_c^{(\mathrm{F})}}{\left(\varepsilon_a^{(\mathrm{Y})} - \varepsilon_c^{(\mathrm{Y})}\right)^3} + \frac{1}{\left(\varepsilon_a^{(\mathrm{Y})} - \varepsilon_c^{(\mathrm{Y})}\right)^2}\left(\frac{\partial \sigma_a^{(\mathrm{F})}}{\partial \varepsilon_c^{(\mathrm{Y})}} + \frac{\partial \sigma_c^{(\mathrm{F})}}{\partial \varepsilon_a^{(\mathrm{Y})}} - \frac{\partial \sigma_a^{(\mathrm{F})}}{\partial \varepsilon_a^{(\mathrm{Y})}} - \frac{\partial \sigma_c^{(\mathrm{F})}}{\partial \varepsilon_c^{(\mathrm{Y})}}\right) \tag{S1.38c}$$

$$\begin{aligned} s_4 = 2\varepsilon_c^{(\mathrm{Y})} \frac{\sigma_a^{(\mathrm{F})} - \sigma_c^{(\mathrm{F})}}{\left(\varepsilon_a^{(\mathrm{Y})} - \varepsilon_c^{(\mathrm{Y})}\right)^3} &+ \frac{1}{\varepsilon_a^{(\mathrm{Y})} - \varepsilon_c^{(\mathrm{Y})}}\left(\frac{\partial \sigma_a^{(\mathrm{F})}}{\partial \varepsilon_c^{(\mathrm{Y})}} - \frac{\partial \sigma_c^{(\mathrm{F})}}{\partial \varepsilon_b^{(\mathrm{Y})}}\right) \\ &+ \frac{\varepsilon_c^{(\mathrm{Y})}}{\left(\varepsilon_a^{(\mathrm{Y})} - \varepsilon_c^{(\mathrm{Y})}\right)^2}\left(\frac{\partial \sigma_a^{(\mathrm{F})}}{\partial \varepsilon_c^{(\mathrm{Y})}} + \frac{\partial \sigma_c^{(\mathrm{F})}}{\partial \varepsilon_a^{(\mathrm{Y})}} - \frac{\partial \sigma_a^{(\mathrm{F})}}{\partial \varepsilon_a^{(\mathrm{Y})}} - \frac{\partial \sigma_c^{(\mathrm{F})}}{\partial \varepsilon_c^{(\mathrm{Y})}}\right) \end{aligned} \tag{S1.38d}$$

$$\begin{aligned} s_5 = 2\varepsilon_c^{(\mathrm{Y})} \frac{\sigma_a^{(\mathrm{F})} - \sigma_c^{(\mathrm{F})}}{\left(\varepsilon_a^{(\mathrm{Y})} - \varepsilon_c^{(\mathrm{Y})}\right)^3} &+ \frac{1}{\varepsilon_a^{(\mathrm{Y})} - \varepsilon_c^{(\mathrm{Y})}}\left(\frac{\partial \sigma_c^{(\mathrm{F})}}{\partial \varepsilon_a^{(\mathrm{Y})}} - \frac{\partial \sigma_c^{(\mathrm{F})}}{\partial \varepsilon_b^{(\mathrm{Y})}}\right) \\ &+ \frac{\varepsilon_c^{(\mathrm{Y})}}{\left(\varepsilon_a^{(\mathrm{Y})} - \varepsilon_c^{(\mathrm{Y})}\right)^2}\left(\frac{\partial \sigma_a^{(\mathrm{F})}}{\partial \varepsilon_c^{(\mathrm{Y})}} + \frac{\partial \sigma_c^{(\mathrm{F})}}{\partial \varepsilon_a^{(\mathrm{Y})}} - \frac{\partial \sigma_a^{(\mathrm{F})}}{\partial \varepsilon_a^{(\mathrm{Y})}} - \frac{\partial \sigma_c^{(\mathrm{F})}}{\partial \varepsilon_c^{(\mathrm{Y})}}\right) \end{aligned} \tag{S1.38e}$$

$$\begin{aligned} s_6 = 2\varepsilon_c^{(\mathrm{Y})} \frac{\sigma_a^{(\mathrm{F})} - \sigma_c^{(\mathrm{F})}}{\left(\varepsilon_a^{(\mathrm{Y})} - \varepsilon_c^{(\mathrm{Y})}\right)^3} &+ \frac{\varepsilon_a^{(\mathrm{Y})}\varepsilon_c^{(\mathrm{Y})}}{\left(\varepsilon_a^{(\mathrm{Y})} - \varepsilon_c^{(\mathrm{Y})}\right)^2}\left(\frac{\partial \sigma_a^{(\mathrm{F})}}{\partial \varepsilon_c^{(\mathrm{Y})}} + \frac{\partial \sigma_c^{(\mathrm{F})}}{\partial \varepsilon_a^{(\mathrm{Y})}}\right) - \frac{\left(\varepsilon_c^{(\mathrm{Y})}\right)^2}{\left(\varepsilon_a^{(\mathrm{Y})} - \varepsilon_c^{(\mathrm{Y})}\right)^2}\left(\frac{\partial \sigma_a^{(\mathrm{F})}}{\partial \varepsilon_a^{(\mathrm{Y})}} + \frac{\partial \sigma_c^{(\mathrm{F})}}{\partial \varepsilon_c^{(\mathrm{Y})}}\right) \\ &- \frac{\varepsilon_a^{(\mathrm{Y})} + \varepsilon_c^{(\mathrm{Y})}}{\varepsilon_a^{(\mathrm{Y})} - \varepsilon_c^{(\mathrm{Y})}}\frac{\partial \sigma_c^{(\mathrm{F})}}{\partial \varepsilon_b^{(\mathrm{Y})}} \end{aligned} \tag{S1.38f}$$

are constants with $(a, b, c)$ being cyclic permutations of (1,2,3).

In the third case, the three eigenvalues of $\varepsilon_{ij}^{(\mathrm{Y})}$ are all identical ($\varepsilon_1^{(\mathrm{Y})} = \varepsilon_2^{(\mathrm{Y})} = \varepsilon_3^{(\mathrm{Y})}$) which gives the following expression for $\mathbf{C}^{(\mathrm{F})}$

$$\mathbf{C}^{(\mathrm{F})} = \left(\frac{\partial \sigma_1^{(\mathrm{F})}}{\partial \varepsilon_1^{(\mathrm{Y})}} - \frac{\partial \sigma_1^{(\mathrm{F})}}{\partial \varepsilon_2^{(\mathrm{Y})}}\right)\mathbf{I}_\mathrm{S} + \frac{\partial \sigma_1^{(\mathrm{F})}}{\partial \varepsilon_2^{(\mathrm{Y})}}\mathbf{I} \otimes \mathbf{I} \tag{S1.39}$$

Note that we need $\partial\sigma_i^{(\mathrm{F})} / \partial\varepsilon_j^{(\mathrm{Y})}$ for all three cases which can be determined by taking the first derivative of $\sigma_i^{(\mathrm{F})}$ in (S1.30)

$$\frac{\partial\sigma_i^{(\mathrm{F})}}{\partial\varepsilon_i^{(\mathrm{Y})}} = \frac{\partial}{\partial\varepsilon_i^{(\mathrm{Y})}}\left(\frac{\partial f}{\partial\varepsilon_i^{(\mathrm{Y})}}\right) = \begin{cases} 0 & \varepsilon_i^{(\mathrm{Y})} < \epsilon_1 \\ \ell\dfrac{\left(\dfrac{\varepsilon_i^{(\mathrm{Y})} - \epsilon_1}{\epsilon_2 - \epsilon_1}\right)^t \left(\varepsilon_i^{(\mathrm{Y})} - \epsilon_1\right)}{t+1} & \epsilon_1 \le \varepsilon_i^{(\mathrm{Y})} < \epsilon_2 \\ \ell\left[\dfrac{\left(1 + \varepsilon_i^{(\mathrm{Y})} - \epsilon_2\right)^{s+1} - 1}{s+1} + \dfrac{\epsilon_2 - \epsilon_1}{t+1}\right] & \varepsilon_i^{(\mathrm{Y})} \ge \epsilon_2 \end{cases}$$

(S1.40)

With $\mathbf{C}^{(\mathrm{F})}$ at hand, the stiffness of the actin filament network $\mathbf{C}^{(\mathrm{A})}$ can be obtained from equation (S1.21).

##### 1.2. Total cell stiffness

To determine the total stiffness of the cell, we first degrade the fourth-order tensors $\mathbf{C}^{(\mathrm{MT})}$ (S1.16), $\mathbf{C}^{(\rho)}$ (S1.19), and $\mathbf{C}^{(\mathrm{A})}$ (S1.21) to the second-order tensors $\mathbf{C}^{(\mathrm{MT})}$, $\mathbf{C}^{(\rho)}$, and $\mathbf{C}^{(\mathrm{A})}$. In Equation (1.20) we show how a fourth-order tensor (*e.g.*, $\mathrm{C}_{ijkl}^{(\rho)}$) is degraded to a second-order 6×6 matrix (*e.g.*, $\mathrm{C}_{ij}^{(\rho)}$).

Note that the microtubule network is connected to the myosin in parallel, and they are both connected to the actin network in series (Fig. S7). Therefore, the total stiffness of the cell, $\mathbf{C}$, is obtained as follows

$$\mathbf{C} = \left(\left(\mathbf{C}^{(\mathrm{X})}\right)^{-1} + \left(\mathbf{C}^{(\mathrm{Y})}\right)^{-1}\right)^{-1} \tag{S1.41}$$

where

$$\mathbf{C}^{(\mathrm{X})} = \mathbf{C}^{(\rho)} + \mathbf{C}^{(\mathrm{MT})} \tag{S1.42}$$

and

$$\mathbf{C}^{(\mathrm{Y})} = \mathbf{C}^{(\mathrm{A})} = \mathbf{C}^{(\mathrm{I})} + \mathbf{C}^{(\mathrm{F})} \tag{S1.43}$$

##### 1.3. Solving the set of nonlinear equations

In the previous sections, we presented the constitutive equations for the cytoskeletal model where the stress field $\sigma_{ij}$ is obtained from equation (S1.17) or (S1.25) and the stiffness field $\mathrm{C}_{ij}$ is determined from (S1.41). However, note that the stress and stiffness tensors $\sigma_{ij}$ and $\mathrm{C}_{ij}$ are functions of the unknown strain tensors $\varepsilon_{ij}^{(\mathrm{X})}$ and $\varepsilon_{ij}^{(\mathrm{Y})}$ (and not $\varepsilon_{ij}$). To determine $\varepsilon_{ij}^{(\mathrm{X})}$ and $\varepsilon_{ij}^{(\mathrm{Y})}$, we first define the following 12×1 vector

$$\boldsymbol{u} = \left\{\varepsilon_{11}^{(X)} \quad \varepsilon_{22}^{(X)} \quad \varepsilon_{33}^{(X)} \quad \varepsilon_{12}^{(X)} \quad \varepsilon_{13}^{(X)} \quad \varepsilon_{23}^{(X)} \quad \varepsilon_{11}^{(Y)} \quad \varepsilon_{22}^{(Y)} \quad \varepsilon_{33}^{(Y)} \quad \varepsilon_{12}^{(Y)} \quad \varepsilon_{13}^{(Y)} \quad \varepsilon_{23}^{(Y)}\right\}^{\mathrm{T}} = \{u_1 \quad u_2 \quad \cdots \quad u_{12}\}^{\mathrm{T}} \tag{S1.44}$$

which contains all 12 unknown variables in the strain tensors $\varepsilon_{ij}^{(X)}$ and $\varepsilon_{ij}^{(Y)}$. To determine the 12 unknowns, we need 12 equations. As the actin filament is connected to other elements in series (Fig. S7), we use the following condition which gives us 6 equations

$$\boldsymbol{\sigma} = \boldsymbol{\sigma}^{(X)} = \boldsymbol{\sigma}^{(Y)} \tag{S1.45}$$

where the stress $\boldsymbol{\sigma}^{(X)}$ (equations (S1.17) and (S1.18))

$$\sigma_{ij}^{(X)} = \sigma_{ij} = \left(C_{ijkl}^{(\rho)} + C_{ijkl}^{(MT)}\right)\varepsilon_{kl}^{(X)} + \bar{\rho}_0\delta_{ij} \tag{S1.46}$$

is directly transmitted to the actin filament network $\boldsymbol{\sigma}^{(Y)}$ (equation (S1.41))

$$\sigma_{ij}^{(Y)} = \sigma_{ij} = \sigma_{ij}^{(A)} = \sigma_{ij}^{(I)} + \sigma_{ij}^{(F)} \tag{S1.47}$$

We get the other 6 equations from the following condition

$$\boldsymbol{\varepsilon} = \boldsymbol{\varepsilon}^{(X)} + \boldsymbol{\varepsilon}^{(Y)} \tag{S1.48}$$

where $\boldsymbol{\varepsilon}^{(X)}$ is the strain of the cytoskeletal components that are in compression (*e.g.*, microtubule network), $\boldsymbol{\varepsilon}^{(Y)}$ is the strain of the cytoskeletal components that are in tension (*e.g.*, actin filaments), and $\boldsymbol{\varepsilon}$ is the total strain of the cell (Fig. S7). Note that all stress and strain tensors $\sigma_{ij}^{(X)}$, $\sigma_{ij}^{(Y)}$, $\varepsilon_{ij}^{(X)}$, and $\varepsilon_{ij}^{(Y)}$ are symmetric. Therefore, the conditions in (S1.45) and (S1.48) can be defined by the following 12 equations in the 12×1 vector $\boldsymbol{f}$

$$\boldsymbol{f} = \{f_1 \quad f_2 \quad \cdots \quad f_{12}\}^{\mathrm{T}} \tag{S1.49a}$$

where

$$f_1 = \sigma_{11}^{(X)} - \sigma_{11}^{(Y)} \tag{S1.49b}$$

$$f_2 = \sigma_{22}^{(X)} - \sigma_{22}^{(Y)} \tag{S1.49c}$$

$$f_3 = \sigma_{33}^{(X)} - \sigma_{33}^{(Y)} \tag{S1.49d}$$

$$f_4 = \sigma_{12}^{(X)} - \sigma_{12}^{(Y)} \tag{S1.49e}$$

$$f_5 = \sigma_{13}^{(X)} - \sigma_{13}^{(Y)} \tag{S1.49f}$$

$$f_6 = \sigma_{23}^{(X)} - \sigma_{23}^{(Y)} \tag{S1.49g}$$

$$f_7 = \varepsilon_{11} - \varepsilon_{11}^{(X)} - \varepsilon_{11}^{(Y)} \tag{S1.49h}$$

$$f_8 = \varepsilon_{22} - \varepsilon_{22}^{(X)} - \varepsilon_{22}^{(Y)} \tag{S1.49i}$$

$$f_9 = \varepsilon_{33} - \varepsilon_{33}^{(X)} - \varepsilon_{33}^{(Y)} \tag{S1.49j}$$

$$f_{10} = \varepsilon_{12} - \varepsilon_{12}^{(X)} - \varepsilon_{12}^{(Y)} \tag{S1.49k}$$

$$f_{11} = \varepsilon_{13} - \varepsilon_{13}^{(X)} - \varepsilon_{13}^{(Y)} \tag{S1.49l}$$

$$f_{12} = \varepsilon_{23} - \varepsilon_{23}^{(X)} - \varepsilon_{23}^{(Y)} \tag{S1.49m}$$

We then determine the 12×12 Jacobian matrix $\mathbf{J}$

$$\mathbf{J} = \begin{bmatrix} \partial f_1/\partial u_1 & \partial f_1/\partial u_2 & \dots & \partial f_1/\partial u_{12} \\ \partial f_2/\partial u_1 & \partial f_2/\partial u_2 & \dots & \partial f_2/\partial u_{12} \\ \vdots & \vdots & & \vdots \\ \partial f_{12}/\partial u_1 & \partial f_{12}/\partial u_2 & \dots & \partial f_{12}/\partial u_{12} \end{bmatrix} = \begin{bmatrix} \mathbf{C}^{(X)} & -\mathbf{C}^{(Y)} \\ -\mathbf{I} & -\mathbf{I} \end{bmatrix} \tag{S1.50}$$

using equations (S1.44) and (S1.49) for $u_i$ and $f_i$, respectively. Finally, we use the Newton-Raphson method

$$\boldsymbol{u}_{i+1} = \boldsymbol{u}_i - \mathbf{J}^{-1}\boldsymbol{f}(\boldsymbol{u}_i) \tag{S1.51}$$

to determine the unknown vector $\boldsymbol{u}$**,** where the vectors $\boldsymbol{u}_i$ and $\boldsymbol{u}_{i+1}$ are respectively the solutions for $i$ and $i+1$ iterations. To find the solutions to the nonlinear equations, we use the following initial guess $\boldsymbol{u}_0$ (or any other initial guess)

$$\boldsymbol{u}_0 = \{0 \quad 0 \quad \dots \quad 0\}^{\mathrm{T}}_{1\times 12} \tag{S1.52}$$

and the convergence criterion

$$|\boldsymbol{f}| = \sqrt{(f_1)^2 + (f_2)^2 + \cdots + (f_{12})^2} < \epsilon_{\mathrm{Tol}} \tag{S1.53}$$

where $|\boldsymbol{f}|$ is the magnitude of the vector $\boldsymbol{f}$, and $\epsilon_{\mathrm{Tol}}$ is the convergence threshold. Using $\epsilon_{\mathrm{Tol}} = 10^{-8}$ in our simulations, we stop the iterations in (S1.51) when $|\boldsymbol{f}|$ is less than $\epsilon_{\mathrm{Tol}}$. With the strain tensors $\varepsilon_{ij}^{(X)}$ and $\varepsilon_{ij}^{(Y)}$ determined from (S1.51), we can calculate the stress tensor $\sigma_{ij}$ (from (S1.17) or (S1.25)) and the stiffness tensor $\mathrm{C}_{ij}$ (from (S1.41)).

##### 1.4. Actomyosin contractility increases with anisotropy in tension

Combining equation (S1.17) with (S1.18), one can derive the following expression for the average of contractility, $\frac{1}{3}\rho_{kk} = (\rho_{11} + \rho_{22} + \rho_{33})/3$

$$\frac{\rho_{kk}}{3} = \left(\frac{3K^{(\mathrm{MT})}\alpha_{\mathrm{v}} - 1}{3K^{(\mathrm{MT})}\beta_{\mathrm{v}} - 1}\right)\frac{\sigma_{kk}}{3} + \left(\frac{3K^{(\mathrm{MT})}\beta_{\mathrm{v}}}{3K^{(\mathrm{MT})}\beta_{\mathrm{v}} - 1}\right)\rho_{0}, \tag{S1.54}$$

which is an expanded form of equation (S1.9) and shows that the average of contractility increases with the average of stress, $\frac{1}{3}\sigma_{kk} = (\sigma_{11} + \sigma_{22} + \sigma_{33})/3$.

We have previously shown that the feedback mechanism between $\rho_{kk}$ and $\sigma_{kk}$ in (S1.54) predicts that contractility, cytoskeleton tension, and traction forces increase with substrate stiffness and substrate area, which are consistent with experimental observations.[5] However, we have recently found that, in addition to the magnitude of tension, cells also respond to the anisotropy of tension.[9] The model was expanded to account for the effect of tension anisotropy where phosphorylation of myosin increases with anisotropy in the components of the stress tensor $\sigma_{ij}$. The increase in actomyosin contractility in response to tension anisotropy is introduced to the model with the term $\alpha_{\mathrm{a}}\sigma_{\mathrm{a}}$

$$\frac{\rho_{kk}}{3} = \left(\frac{3K^{(\mathrm{MT})}\alpha_{\mathrm{v}} - 1}{3K^{(\mathrm{MT})}\beta_{\mathrm{v}} - 1}\right)\frac{\sigma_{kk}}{3} + \alpha_{\mathrm{a}}\sigma_{\mathrm{a}} + \left(\frac{3K^{(\mathrm{MT})}\beta_{\mathrm{v}}}{3K^{(\mathrm{MT})}\beta_{\mathrm{v}} - 1}\right)\rho_{0} \tag{S1.55}$$

where $\sigma_1 > \sigma_2 > \sigma_3$ are the principal stress values (eigenvalues) of the stress tensor $\sigma_{ij}$ with $\sigma_1 > \sigma_2 > 0$, $\sigma_{\mathrm{a}} = \tanh\left(\frac{1}{2}\left(\frac{\sigma_1}{\sigma_2} - 1\right)\right)\rho_0$ represents the tension anisotropy, $\alpha_{\mathrm{a}}$ is the anisotropic chemo-mechanical feedback parameter which regulates the increase in myosin phosphorylation with tension anisotropy. The additional term $\alpha_{\mathrm{a}}\sigma_{\mathrm{a}}$ is implemented into the finite element framework using a piecewise linear approximation where $\rho_0$ in equation (S1.13) is simply replaced with $\rho_0 + \left(\frac{3K^{(\mathrm{MT})}\beta_{\mathrm{v}}-1}{3K^{(\mathrm{MT})}\beta_{\mathrm{v}}}\right)\alpha_{\mathrm{a}}\sigma_{\mathrm{a}}$ in each step of the simulation.

#### 2. Total free energy

We hypothesized that the phenotypic transitions observed in cells, whether adopting a round morphology, forming spherical clusters, or elongating, can be predicted by minimizing the system's total free energy. Following the second law of thermodynamics, all processes, including the cell's interactions with the ECM, evolve toward configurations that reduce the overall free energy of the system. Here, the total free energy is defined as the sum of the free energy of the cell and that of the ECM, reflecting the coupled mechanical and chemical contributions from both intracellular and extracellular components.

##### 2.1. Cell energy

We begin by evaluating the energetic contributions within our cytoskeletal model. The cell's total free energy is decomposed into three distinct components: motor-work energy, chemical energy, and mechanical energy.[1]

The motor-work energy arises from the active contractile behavior of myosin molecular motors, which convert chemical energy derived from ATP hydrolysis into mechanical work:

$$\mathrm{W}_{\text{motor-work}} = \frac{1}{3}\,\rho_{kk}\,\varepsilon_{kk}^{(\mathrm{X})} + \tilde{\rho}_{ij}\,\tilde{\varepsilon}_{ij}^{(\mathrm{X})} \tag{S2.1}$$

where $\tilde{\rho}_{ij}$ and $\tilde{\varepsilon}_{ij}^{(\mathrm{X})}$ denote the deviatoric parts of the contractility tensor and the strain tensor (in the contractile unit), respectively, and are defined as:

$$\begin{aligned} \tilde{\rho}_{ij} &= \rho_{ij} - \frac{1}{3}\rho_{kk}\delta_{ij} \\ \tilde{\varepsilon}_{ij}^{(\mathrm{X})} &= \varepsilon_{ij}^{(\mathrm{X})} - \frac{1}{3}\varepsilon_{kk}^{(\mathrm{X})}\delta_{ij} \end{aligned} \tag{S2.2}$$

The chemical energy term captures the regulation of myosin motor phosphorylation in response to cytoskeletal tension. It reflects a balance between the energetic cost of recruiting myosin motors and the chemical energy released through their phosphorylation:

$$\mathrm{W}_{\text{chemical}} = \frac{\beta_{\mathrm{v}}}{2}\frac{1}{3}(\rho_{kk} - 3\rho_0)^2 + \frac{\beta_{\mathrm{d}}}{2}\tilde{\rho}_{ij}\tilde{\rho}_{ij} - \frac{1}{3}\int_0^{\rho_{kk}} \alpha_{\mathrm{v}}\sigma_{kk}\,d\rho_{kk} - \int_0^{\tilde{\rho}_{ij}} \alpha_{\mathrm{d}}\tilde{\sigma}_{ij}\,d\tilde{\rho}_{ij} \tag{S2.3}$$

where $\tilde{\sigma}_{ij}$ denotes the deviatoric part of the stress tensor, and is defined as:

$$\tilde{\sigma}_{ij} = \sigma_{ij} - \frac{1}{3}\sigma_{kk}\delta_{ij} \tag{S2.4}$$

In the absence of external stress, the motor density stabilizes at $\rho_0$, a value determined by the equilibrium between myosin binding and unbinding events, thus minimizing the chemical free energy. Deviations from this equilibrium due to increases in cytoskeletal tension, activate signaling pathways such as $Ca^{2+}$ influx or Rho-ROCK, leading to enhanced myosin recruitment. In this formulation, the first two terms penalize deviations from $\rho_0$, while the second two terms represent chemo-mechanical coupling, where mechanical stress facilitates contractility and reduces the system's free energy.

The mechanical energy captures the elastic response of the cytoskeleton, particularly the deformation of passive structural elements such as microtubules and actin filaments. This energy contribution reflects the resistance of the cytoskeletal framework to applied strain and is written as:

$$\mathrm{W}_{\mathrm{mechanical}} = \frac{K^{(\mathrm{MT})}}{2}(\varepsilon_{kk}^{(\mathrm{X})})^2 + \mu^{(\mathrm{MT})}\tilde{\varepsilon}_{ij}^{(\mathrm{X})}\tilde{\varepsilon}_{ij}^{(\mathrm{X})} - \frac{1}{3}\int_0^{\varepsilon_{kk}^{(\mathrm{X})}} \sigma_{kk} d\varepsilon_{kk}^{(\mathrm{X})} - \int_0^{\tilde{\varepsilon}_{ij}^{(\mathrm{X})}} \tilde{\sigma}_{ij} d\tilde{\varepsilon}_{ij}^{(\mathrm{X})} + \mathrm{W}_{\mathrm{actin}} \tag{S2.5}$$

To calculate the energy contribution from the actin network, we note that the actin element is modeled as a nonlinear strain-stiffening material. Accordingly, the actin energy can be expressed as:

$$\mathrm{W}_{\mathrm{actin}} = \mathrm{W}_{\mathrm{actin}}^{(\mathrm{I})} + \mathrm{W}_{\mathrm{actin}}^{(\mathrm{F})} \tag{S2.6}$$

$\mathrm{W}_{\mathrm{actin}}^{(\mathrm{I})}$ accounts for the isotropic linear part of the actin stiffness:

$$\mathrm{W}_{\mathrm{actin}}^{(\mathrm{I})} = \frac{K^{(\mathrm{A})}}{2}(\varepsilon_{kk}^{(\mathrm{Y})})^2 + \mu^{(\mathrm{A})}\tilde{\varepsilon}_{ij}^{(\mathrm{Y})}\tilde{\varepsilon}_{ij}^{(\mathrm{Y})} \tag{S2.7}$$

where $K^{(\mathrm{A})}$ and $\mu^{(\mathrm{A})}$ are the bulk and shear moduli of the actin network, respectively, and are defined as:

$$K^{(\mathrm{A})} = \frac{E^{(\mathrm{A})}}{3(1 - 2v^{(\mathrm{A})})}$$
$$\mu^{(\mathrm{A})} = \frac{E^{(\mathrm{A})}}{2(1 + v^{(\mathrm{A})})} \tag{S2.8}$$

Here, $E^{(\mathrm{A})}$ and $v^{(\mathrm{A})}$ are the elastic modulus and the Poisson's ratio of the actin network. $\tilde{\varepsilon}_{ij}^{(\mathrm{Y})}$ in (S2.7) denotes the deviatoric part of the strain tensor in the actin network, and is defined as:

$$\tilde{\varepsilon}_{ij}^{(\mathrm{Y})} = \varepsilon_{ij}^{(\mathrm{Y})} - \frac{1}{3}\varepsilon_{kk}^{(\mathrm{Y})}\delta_{ij} \tag{S2.9}$$

To account for the nonlinear contribution of the actin energy, we note that $\sigma_i^{(\mathrm{F})}$ in (S1.29) is the derivative of the nonlinear component of actin energy with respect to strain. Therefore, the nonlinear actin energy can be obtained by integrating equation (S1.29) with respect to principal strains $\varepsilon_i^{(\mathrm{Y})}$:

$$
\mathrm{W}_{\text{actin}}^{(\mathrm{F})} = \begin{cases} 0 & \varepsilon_i^{(\mathrm{Y})} < \epsilon_1 \\ \ell \dfrac{\left(\dfrac{\varepsilon_i^{(\mathrm{Y})} - \epsilon_1}{\epsilon_2 - \epsilon_1}\right)^{t+3} (\epsilon_2 - \epsilon_1)^3}{(t+1)(t+2)(t+3)} & \epsilon_1 \le \varepsilon_i^{(\mathrm{Y})} < \epsilon_2 \\ \ell \left[ \dfrac{\left(1 + \varepsilon_i^{(\mathrm{Y})} - \epsilon_2\right)^{s+3}}{(s+1)(s+2)(s+3)} + \dfrac{\epsilon_2 \varepsilon_i^{(\mathrm{Y})} - 0.5\left(\varepsilon_i^{(\mathrm{Y})}\right)^2}{s+1} - \dfrac{\varepsilon_i^{(\mathrm{Y})}}{(s+1)(s+2)} + \dfrac{0.5(\varepsilon_i^{(\mathrm{Y})} - \epsilon_2)^2(\epsilon_2 - \epsilon_1)}{t+1} + \dfrac{(\epsilon_2 - \epsilon_1)^2 \varepsilon_i^{(\mathrm{Y})}}{(t+1)(t+2)} \right] & \varepsilon_i^{(\mathrm{Y})} \ge \epsilon_2 \end{cases} \quad \text{(S2.10)}
$$

Together, these energy terms define the total free energy of the cell in our three-dimensional model.

#### 2.2. ECM energy

In addition to the cell energy terms, deformation of the ECM contributes to the overall energy of the system. ECM energy can be expressed as:

$$
\mathrm{W}_{\mathrm{ECM}} = \mathrm{W}_{\mathrm{ECM}}^{\text{expansion}} + \mathrm{W}_{\mathrm{ECM}}^{\text{contraction}} \quad \text{(S2.11)}
$$

where $\mathrm{W}_{\mathrm{ECM}}^{\text{expansion}}$ and $\mathrm{W}_{\mathrm{ECM}}^{\text{contraction}}$ are energy stored in the ECM due to cell expansion and cell contraction, respectively, and both are formulated as:

$$
\mathrm{W}_{\mathrm{ECM}} = \frac{K^{(\mathrm{ECM})}}{2} (\varepsilon_{kk}^{(\mathrm{ECM})})^2 + \mu^{(\mathrm{ECM})} \tilde{\varepsilon}_{ij}^{(\mathrm{ECM})} \tilde{\varepsilon}_{ij}^{(\mathrm{ECM})} \quad \text{(S2.12)}
$$

where $K^{(\mathrm{ECM})}$ and $\mu^{(\mathrm{ECM})}$ are the bulk and shear moduli of the ECM, respectively, and are defined as:

$$
K^{(\mathrm{ECM})} = \frac{E^{(\mathrm{ECM})}}{3(1 - 2v^{(\mathrm{ECM})})}
$$

$$
\mu^{(\mathrm{ECM})} = \frac{E^{(\mathrm{ECM})}}{2(1 + v^{(\mathrm{ECM})})} \quad \text{(S2.13)}
$$

Here, $E^{(\mathrm{ECM})}$ and $v^{(\mathrm{ECM})}$ are the elastic modulus and the Poisson's ratio of the ECM, respectively. $\tilde{\varepsilon}_{ij}^{(\mathrm{ECM})}$ in (S2.12) is the deviatoric part of the strain tensor in the ECM, and is defined as:

$$
\tilde{\varepsilon}_{ij}^{(\mathrm{ECM})} = \varepsilon_{ij}^{(\mathrm{ECM})} - \frac{1}{3} \varepsilon_{kk}^{(\mathrm{ECM})} \delta_{ij} \quad \text{(S2.14)}
$$

The total free energy of the system is determined by the sum of the cell free energy and the ECM free energy:

$$W_{\mathrm{total}} = W_{\mathrm{cell}} + W_{\mathrm{ECM}} = W_{\mathrm{motor-work}} + W_{\mathrm{chemical}} + W_{\mathrm{mechanical}} + W_{\mathrm{ECM}} \tag{S2.15}$$

### 3. Simulations

Finite element simulations were performed by implementing the theoretical model detailed in Supplementary Information 1 into a UMAT subroutine within Abaqus. This framework was used to compute the cell-ECM interaction energy, as described in Supplementary Information 2, across a range of ECM stiffnesses and morphological configurations. Specifically, we modeled three scenarios: a single round cell, a spherical cluster, and a single elongated cell, embedded within either a soft or stiff ECM. For each case, we calculated the individual contributions of motor-work, chemical, and mechanical energy within the cell, along with the strain energy stored in the ECM due to matrix deformation. Since the energy terms described in the previous section represent volumetric densities, we multiplied these values by the corresponding volumes of the cell and the ECM in each scenario to obtain the total energy contributions from each component of the system. These simulations allowed us to quantify how total free energy varies with both ECM stiffness and cell morphology. The code implementing the model in the finite element framework is available on GitHub (https://github.com/Farid-Alisafaei/ECM-Mechanics-Regulate-Cancer-Cell-State) and includes annotations describing the model parameters.

##### 3.1 Single round cell

The cell was modeled as a sphere with a diameter of 25 μm (Figure S4), embedded at the center of an ECM domain. The ECM was represented as a cube with a side length of 800 μm, sufficiently large to eliminate any boundary effects on the mechanical response of the cell. In this scenario, the cell does not expand; therefore, the ECM energy was computed solely based on the strain energy generated by cell contraction. A fine mesh density was selected to ensure sufficient resolution for capturing deformation gradients and achieving numerical accuracy throughout the domain. The cell domain was discretized using 68,616 linear tetrahedral elements (C3D4), while the ECM was meshed with a combination of 8,432 C3D4 elements near the cell and 63,992 linear hexahedral elements (C3D8) in regions distant from the cell.

##### 3.2 Spherical cluster

The spherical cluster configuration was modeled as an aggregate of six individual cells (Figure S4), each represented as a sphere with a diameter of 25 μm, arranged symmetrically with marginal overlap to form a compact structure. This cluster was embedded at the center of an ECM domain modeled as a cube with a side length of 800 μm. To compute the energy stored in the ECM due to cluster expansion, a separate simulation was conducted in which the ECM contained a central void equal in size to a single cell. A displacement field was then applied to the void boundary to expand it to the final size of the six-cell aggregate, allowing estimation of the ECM strain energy associated with cell cluster expansion. After calculating the ECM energy resulting from matrix deformation due to cluster expansion driven by cell proliferation, we next calculated the ECM energy associated with cluster contraction. The ECM energy contribution due to cell contraction was determined from a coupled simulation containing both the cluster and the surrounding ECM. The cluster was discretized with 222,545 linear tetrahedral elements (C3D4). The ECM mesh included 39,415 C3D4 elements near the cluster and 62,250 linear hexahedral elements (C3D8) in the outer region.

##### 3.3 Elongated cell

The elongated cell configuration was modeled as a long prolate spheroid (Figure S4), with a minor axis of 25 µm and a major axis of 150 µm, positioned at the center of a surrounding ECM domain. The ECM was again defined as a cube with a side length of 800 µm, consistent with other simulation setups. To isolate the energy contribution from matrix deformation caused by cell elongation, a separate simulation was performed in which the ECM initially included a central void corresponding to a spherical cell. A prescribed displacement was then applied to the void boundary to expand it to the shape and dimensions of the elongated cell, enabling the calculation of the ECM strain energy resulting from cell expansion. The additional ECM energy due to cell contractility was computed in a separate coupled model including both the elongated cell and the ECM. Note that the processes of cell spreading (elongation) and cell contraction occur cyclically over time, driving the transition from a small, round morphology to a fully spread and elongated state, as we have recently demonstrated.[9] However, for simplicity, we simulated each of these two processes separately and calculated the energy associated with each individually. The cell geometry was meshed using 211,039 linear tetrahedral elements (C3D4). The ECM mesh consisted of 32,041 C3D4 elements in proximity to the cell and 67,096 linear hexahedral elements (C3D8) farther from the cell to ensure accurate resolution of localized and bulk matrix deformation.

The material properties assigned to the ECM, along with the model parameters used for the cell chemo-mechanical formulation, are summarized in Table S1.

#### Supplementary Figures

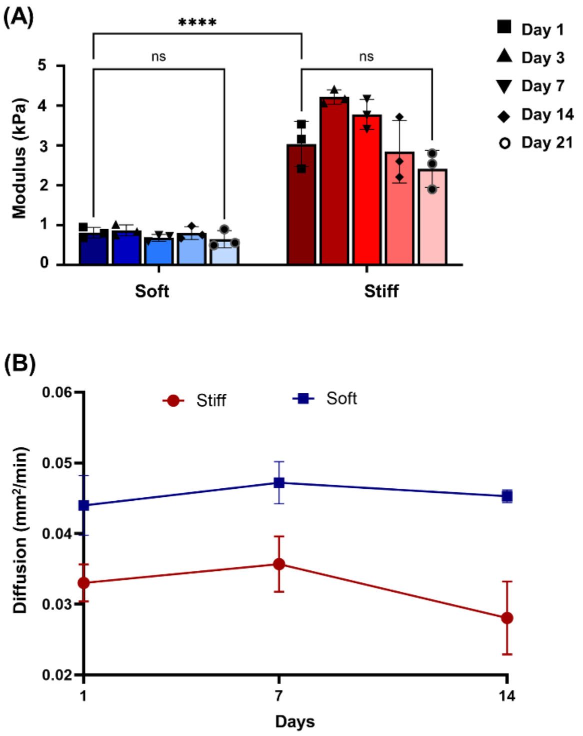


**Figure S1.** (A) Stiffness and (B) diffusion measurements of cell-laden soft and stiff hydrogels over time

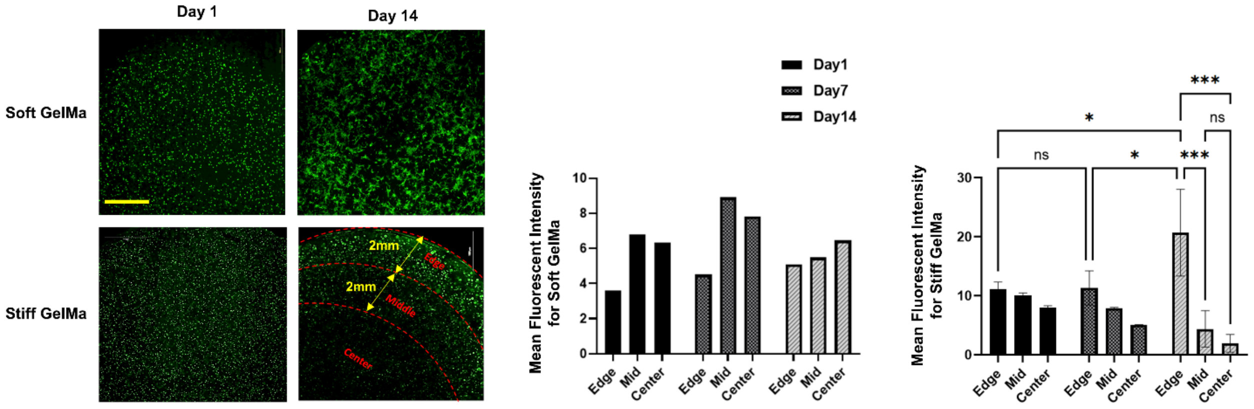


**Figure S2.** Fluorescence images of GFP-labeled MDA-MB-231 cells within soft and stiff hydrogels, acquired at 4× magnification and segmented into center, middle, and edge regions (scale bar: 1000 µm). Unlike cells in soft ECM, cells in stiff ECM showed significantly higher fluorescence intensity at the edge, suggesting that cells near the periphery may be more viable. Accordingly, all measurements in this study were performed using cells located at the edge of the tissues.

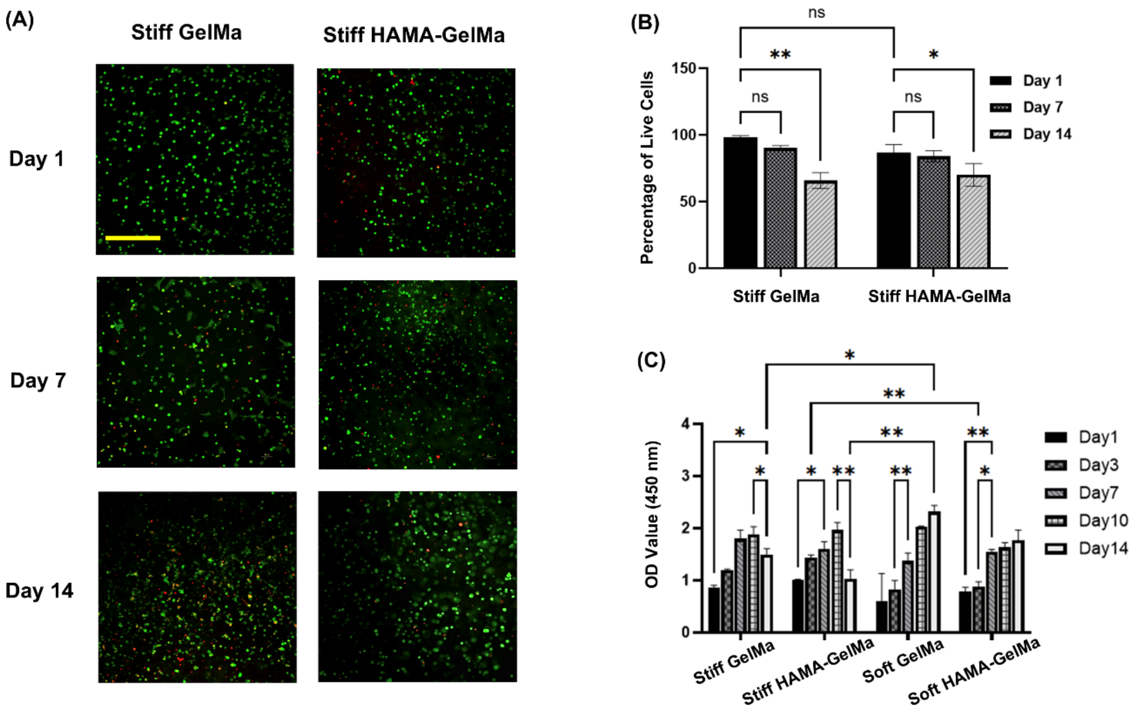


**Figure S3.** (A) Representative images of viability staining in the center of stiff ECMs using Calcein-AM (green, live cells) and propidium iodide (red, dead cells). Scale bar: 100 µm. (B) Quantification of cell viability and (C) metabolic activity over time. Consistent with the spatial distribution observed in Fig. S2, cells in the center of stiff ECMs showed reduced viability and metabolic activity by day 14. This effect was not observed in the center of soft ECMs. Based on these findings, all measurements in this study were performed using cells located at the edge of the tissues.

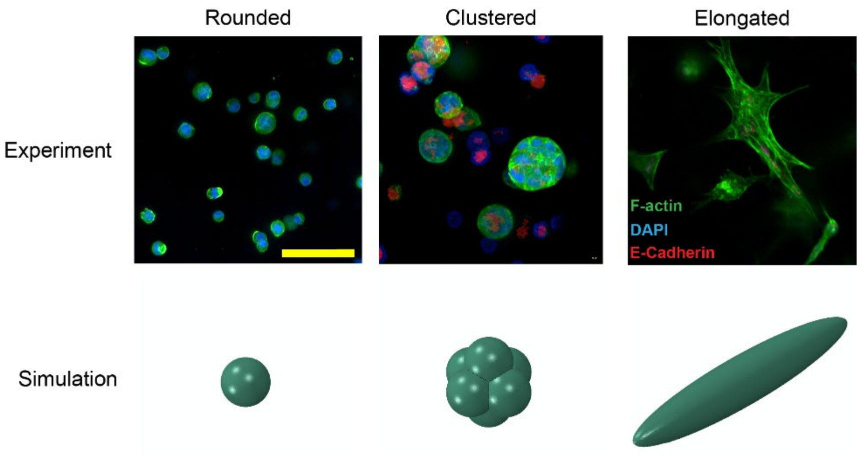


**Figure S4.** Using the theoretical model described in Supplementary Notes 1–3, we simulated the three possible cell morphological states—rounded, elongated, and clustered—within both soft and stiff ECMs. The model calculates the individual energy components for each configuration, including ECM mechanical energy and the three cell energy terms (Fig. 2C), to determine the total energy associated with each morphological state (Fig. 3A).

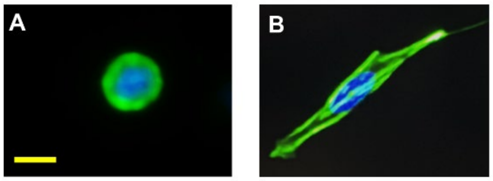


**Figure S5.** Morphological transition of a representative cell within a soft ECM; (A) Initially, the cell exhibits a small, rounded morphology. (B) Over time, the cell spreads and elongates. Scale bar: 10 μm.

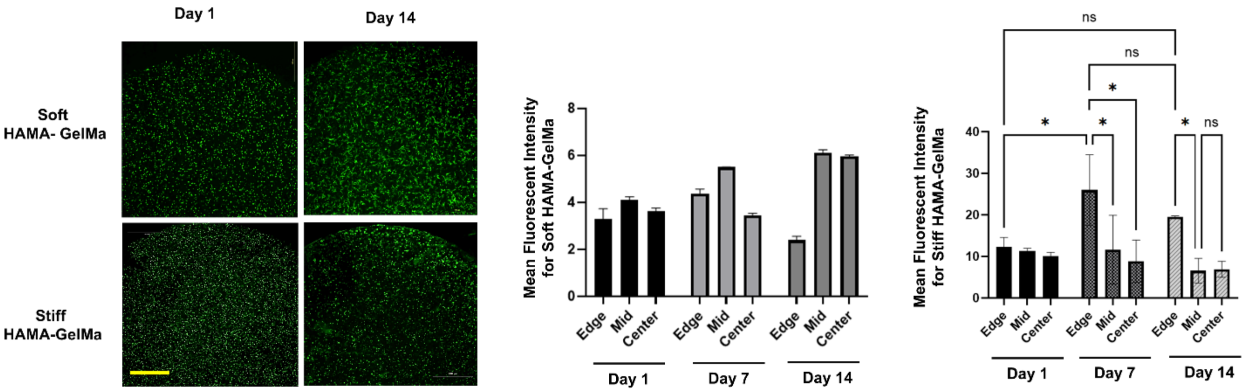


**Figure S6.** Fluorescence images of GFP-expressing MDA-MB-231 cells within soft and stiff HAMA-GelMA hydrogels, acquired at 4× magnification and segmented into center, middle, and edge regions (scale bar: 1000 μm). Similar to the GelMA-only hydrogels shown in Fig. S2, cells in stiff composite ECMs exhibited higher fluorescence intensity at the edges. Quantification of fluorescence intensity across regions is shown for cells both within soft and stiff HAMA-GelMA hydrogels.

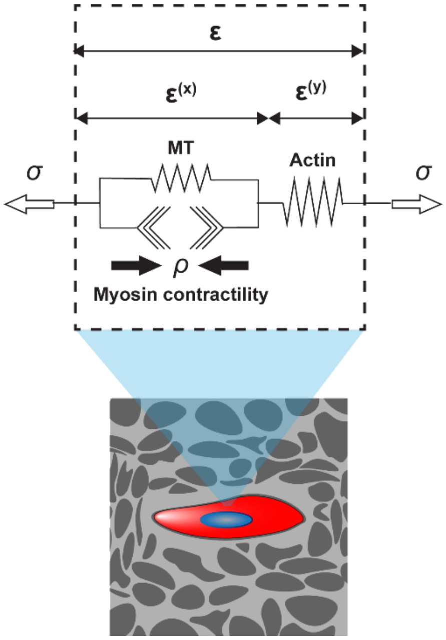


**Figure S7.** 1D representation of $\boldsymbol{\rho}$, $\boldsymbol{\sigma}$, $\boldsymbol{\varepsilon}^{(\mathbf{x})}$, $\boldsymbol{\varepsilon}^{(\mathbf{y})}$, and $\boldsymbol{\varepsilon}$ in our 3D model.

#### Supplementary Tables

**Table S1.** Parameters for the three-dimensional cytoskeletal model.

| Parameter | Description | Value | Unit |
|---|---|---|---|
| $\rho_0$ | Baseline contractility | 16.5 | kPa |
| $\alpha_{\mathrm{v}}$ | Volumetric feedback parameter | 0.3 | 1/kPa |
| $\alpha_{\mathrm{d}}$ | Deviatoric feedback parameter | 0.3 | 1/kPa |
| $\beta_{\mathrm{v}}$ | Volumetric chemical stiffness parameter | 2 | 1/kPa |
| $\beta_{\mathrm{d}}$ | Deviatoric chemical stiffness parameter | 2 | 1/kPa |
| $E^{(\mathrm{MT})}$ | Stiffness of microtubule | 8 | kPa |
| $E^{(\mathrm{A})}$ | Stiffness of actin network | 8 | kPa |
| $\nu^{(\mathrm{MT})}$ | Poisson's ratio of microtubule | 0.4 | |
| $\nu^{(\mathrm{A})}$ | Poisson's ratio of actin | 0.5 | kPa |
| $l$ | Actin network strain-stiffening parameter | 50 | |
| $s$ | Actin network strain-stiffening parameter | 4 | |
| $t$ | Actin network strain-stiffening parameter | 4 | |
| $\epsilon_{\mathrm{c}}$ | Critical tensile principal strain of actin network | 0.1 | |
| $\alpha_{\mathrm{a}}$ | anisotropic chemo-mechanical feedback parameter | 0.4 | |
| ${E^{(\mathrm{ECM})}}_{\mathrm{soft}}$ | Stiffness of soft ECM | 4 | kPa |
| ${\nu^{(\mathrm{ECM})}}_{\mathrm{soft}}$ | Poisson's ratio of soft ECM | 0.3 | |
| ${E^{(\mathrm{ECM})}}_{\mathrm{stiff}}$ | Stiffness of stiff ECM | 1 | kPa |
| ${\nu^{(\mathrm{ECM})}}_{\mathrm{stiff}}$ | Poisson's ratio of stiff ECM | 0.3 | |
